# Modular control of time and space during vertebrate axis segmentation

**DOI:** 10.1101/2023.08.30.555457

**Authors:** Ali Seleit, Ian Brettell, Tomas Fitzgerald, Carina Vibe, Felix Loosli, Joachim Wittbrodt, Kiyoshi Naruse, Ewan Birney, Alexander Aulehla

## Abstract

How temporal and spatial control of developmental processes are linked remains a fundamental question. Do underlying mechanisms form a single functional unit or are these dissociable modules?

We address this question by studying the periodic process of embryonic axis segmentation, using genetic crosses of inbred medaka fish strains representing two species, *Oryzias sakaizumii* and *latipes*. Our analysis revealed correlated interspecies differences with regard to the timing of segmentation, the size of segments and of the presomitic mesoderm (PSM), from which segments are periodically formed. We then did interspecies crosses and real-time imaging quantifications, which revealed extensive phenotypic variation in ∼600 F2 embryos. Importantly, while the F2 analysis showed correlated changes of PSM and segment size, these spatial measures were not correlated to the timing of segmentation. This shows that the control of time and space of axis segmentation can, in principle, be decoupled. In line with this finding, we identified, using *developmental* quantitative trait loci (*dev*QTL) mapping, distinct chromosomal regions linked to either the control of segmentation timing or PSM size. We were able to validate the *dev*QTL findings using a CRISPR/Cas9 loss-of-function approach on several candidate genes *in vivo*.

Combined, this study reveals that a developmental constraint mechanism underlies spatial scaling of axis segmentation, while its spatial and temporal control are dissociable modules. Our findings emphasise the need to reveal the selective constraints linking these modules in the natural environment.

---

How developmental timing is controlled and linked to the form and function of developing organisms remains a fundamental, challenging question. In part, this is due to the fact that timing of development is under complex control, integrating both genetic^1–3^ and environmental factors^4,5^, which combined result in a species-characteristic timing. For instance, in vertebrates, the completion of body axis segmentation into the pre-vertebrae takes ∼15 days in humans^6,7^, ∼4 days in mice^8,9^, ∼3 days in chick^10,11^ and ∼1 day in zebrafish embryos^4,12^. The species-characteristic timing of body axis segmentation is linked to the underlying activity of the segmentation clock^13^. The clock activity can be quantified at the level of ultradian Notch-signalling oscillations that occur in presomitic mesoderm (PSM) cells^14^ with a period matching the species-characteristic rate of addition of somites, ∼6h in human^15,16^, ∼2h in mouse^17^, ∼90 minutes in chick^13^ and ∼30 minutes in zebrafish embryos^18^. How differences in timing relate to distinct morphologies and proportions is challenging to address in evolutionarily distant species^19–21^. We hence employed a comparative, *common garden*^22^ approach using closely related medaka fish species, *Oryzias sakaizumii* and *latipes,* that are native to different regions in Japan. The northern *Oryzias sakaizumii* (Kaga, HNI) and southern *Oryzias latipes* (Cab, HdrII, Ho5) (Fig. 1a) have an estimated evolutionary divergence time of ∼18 million years ^23,24^. Importantly, these species exhibit developmental and phenotypic variation^25–27^, tolerance to inbreeding^27,28^ and are amenable to interbreeding. We developed a real-time imaging approach, which enables a quantitative analysis of both the temporal and spatial measures of embryonic axis segmentation in genetically diverse F2 offsprings resulting from interbreeding of O. *sakaizumii* and *O. latipes.* We combined the phenotypic analysis with whole genome sequencing of ∼600 F2 embryos and performed *developmental* quantitative trait loci (*dev*QTL) mapping to gain insight into how the control of *time* and *space* is functionally linked during development.

**Figure 1:**
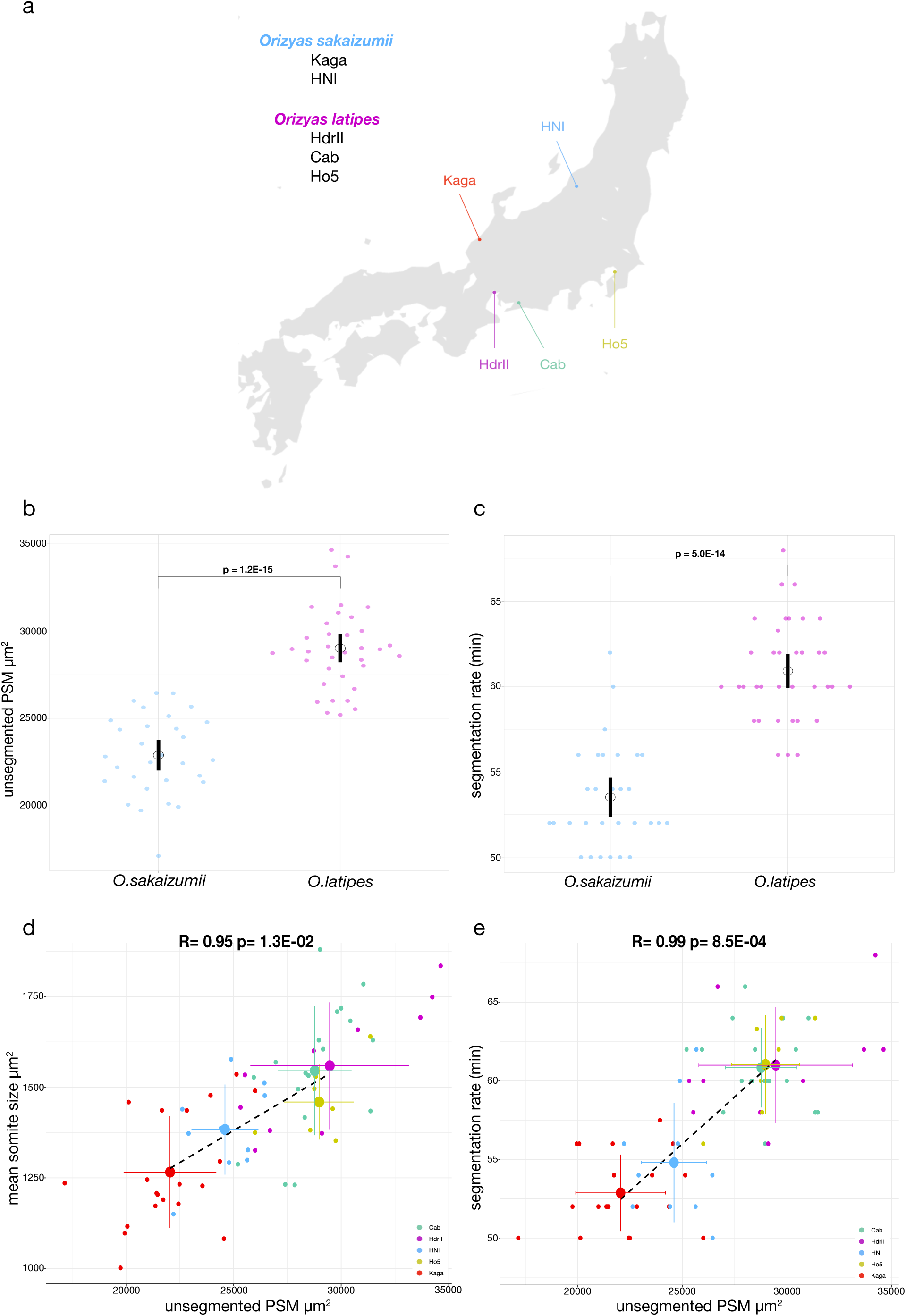
**Scaling of segmentation timing and size in *Oryzias sakaizumii and Oryzias latipes*** (a) Schematic map of mainland Japan (showing Honshu, Shikoku and parts of Kyushu) highlighting the sites of the original medaka populations. The northern *Oryzias sakaizumii*: HNI (blue) and Kaga (red) diverged from the southern *Oryzias latipes*: Cab (turquoise), HdrII (magenta) and Ho5 (yellow) 18 million years ago. (b) unsegmented presomitic mesoderm (PSM) size at the 10-11 somite stage (SS) obtained from brightfield live-imaging*. Oryzias sakaizumii* have a smaller PSM size compared to *Oryzias latipes* (22898μm^2^ (sd +/-2293) vs. 29004μm^2^ (sd +/-2350)). Black circle = mean period. Black line = 95% confidence interval. Welch two sample t-test p= 1.2E-15. N= 30 *Oryzias sakaizumii,* N= 36 *Oryzias latipes.* (c) Embryonic segmentation rate at the 10-11 somite stage (SS) obtained from brightfield live-imaging*. Oryzias sakaizumii* shows a faster segmentation period compared to *Oryzias latipes* (53.52 minutes (sd +/-3.0) *vs*. 60.92 minutes (sd +/-2.91)). Black circle = mean period. Black line = 95% confidence interval. Welch two sample t-test p= 5.0E-14 N= 30 *Oryzias sakaizumii,* N= 36 *Oryzias latipes.* (d) Pearson’s correlation on average trait values of unsegmented presomitic mesoderm (PSM) size and mean nascent somite size at 10-11SS in *Oryzias sakaizumii* Kaga (red), HNI (blue) and *Oryzias latipes* Cab (green), HdrII (magenta), Ho5 (yellow) embryos shows a positive correlation R= 0.95 p-value= 1.3E-02. Black dotted line= linear fit on average trait values, vertical and horizontal lines on both axes represent standard deviation measures. Individual data points are shown for each population N= 19 Cab, N= 10 HdrII, N= 7 Ho5, N= 20 Kaga, N= 10 HNI (e) Pearson’s correlation between average segmentation rate and average unsegmented presomitic mesoderm (PSM) size at the 10-11SS in *Oryzias sakaizumii* Kaga (red), HNI (blue) and *Oryzias latipes* Cab (green), HdrII (magenta), Ho5 (yellow) embryos shows a positive correlation R= 0.99 p-value= 8.5E-04. Black dotted line= linear fit. vertical and horizontal lines on both axes represent standard deviation measures. Individual data points are shown for each population N= 19 Cab, N= 10 HdrII, N= 7 Ho5, N= 20 Kaga, N= 10 HNI.

## Correlation of segmentation timing, PSM and segment size across medaka species

We first performed *common garden* experiments in several inbred representatives of *Oryzias sakaizumii* and *latipes* species to investigate timing and morphology of axis formation and segmentation. We found that *Oryzias sakaizumii* embryos had smaller PSM and nascent somite sizes compared to *Oryzias latipes* embryos (Fig. 1b, Extended Data Fig. 1a-d). To investigate potential differences in developmental timing we measured segment formation rate *in-vivo* in whole embryos. We found that the timing of segment formation differed between *Oryzias sakaizumii* and *Oryzias latipes* embryos, with the former showing a faster segmentation pace (53.52 +/- 3.0 min (mean +/- sd)) as compared to the latter (60.92 +/- 2.91 min (mean +/- sd)) (p=5.0E-14) (Fig. 1c, Extended Data Fig. 1e-g). This timing difference was also present in embryo explants cultured in defined media (Extended Data Fig. 1h-i), indicating it is of intrinsic origin. Importantly, the comparison of timing and size differences across the five inbred populations revealed that the differences are correlated (Fig. 1d-e, Extended Data Fig. 1b-g). At the level of morphology, our results show that embryos with a larger PSM formed proportionally larger segments (Fig. 1d), this correlation was visible across species (Fig. 1d, Extended Data Fig. 1d). In addition, we found a second scaling relation; faster segmentation occurs in embryos of smaller size (Fig. 1e, Extended Data Fig. 1e-g). Taken together these results revealed a correlation of *temporal* and *spatial* measures during the embryonic axis segmentation process. This in turn raised the question of whether the correlations reflect a single developmental process or multiple coupled developmental modules.

## Hybrid F1 embryos show intermediate segmentation timing

Based on our initial findings that revealed differences in timing and morphology between northern and southern species of medaka, we performed inter-breeding experiments to analyse the degree of heritable variation in F1 and F2 offsprings. We first analysed hybrid F1 offsprings from a series of north-south crosses (Fig. 2a). We used *in-vivo* live-imaging to quantify timing of axis segmentation in hybrid F1 embryos at the 10-11 somite stage (SS) (Fig. 2b-e). To enable a precise quantification of timing differences we made use of a fluorescent segmentation clock endogenous knock-in *her7-venus* reporter line that we recently developed^5^ (Fig. 2b-c, Extended Data Fig. 2a-b). Our results revealed that the segmentation clock period differed across the hybrid F1s (Fig. 2d), with the *Oryzias sakaizumii/latipes* hybrid F1 embryos (HNI/Cab, Kaga/Cab) showing a faster segmentation timing than hybrid F1 *Oryzias latipes/latipes* (Ho5/Cab, HdrII/Cab) (Fig. 2d-e, Extended Data Fig. 2c-d). In addition, *Oryzias sakaizumii/latipes* hybrid F1 embryos showed intermediate segmentation clock period values in between the parental F0 lines (Fig. 2d-e, Extended Data Fig. 2c-d). To address the possible impact of maternal effects, we performed reciprocal (north/south) Kaga/Cab F1 crosses. The results showed that while egg size is maternally controlled as expected, neither PSM size nor segmentation clock period measurements showed evidence for a significant maternal effect (Extended Data Fig. 3a-i). Next, we generated F2 offspring by crossing hybrid F1 Kaga/Cab (north/south) fishes, with the goal of producing individuals having a unique genetic composition following meiotic recombination events.

**Figure 2:**
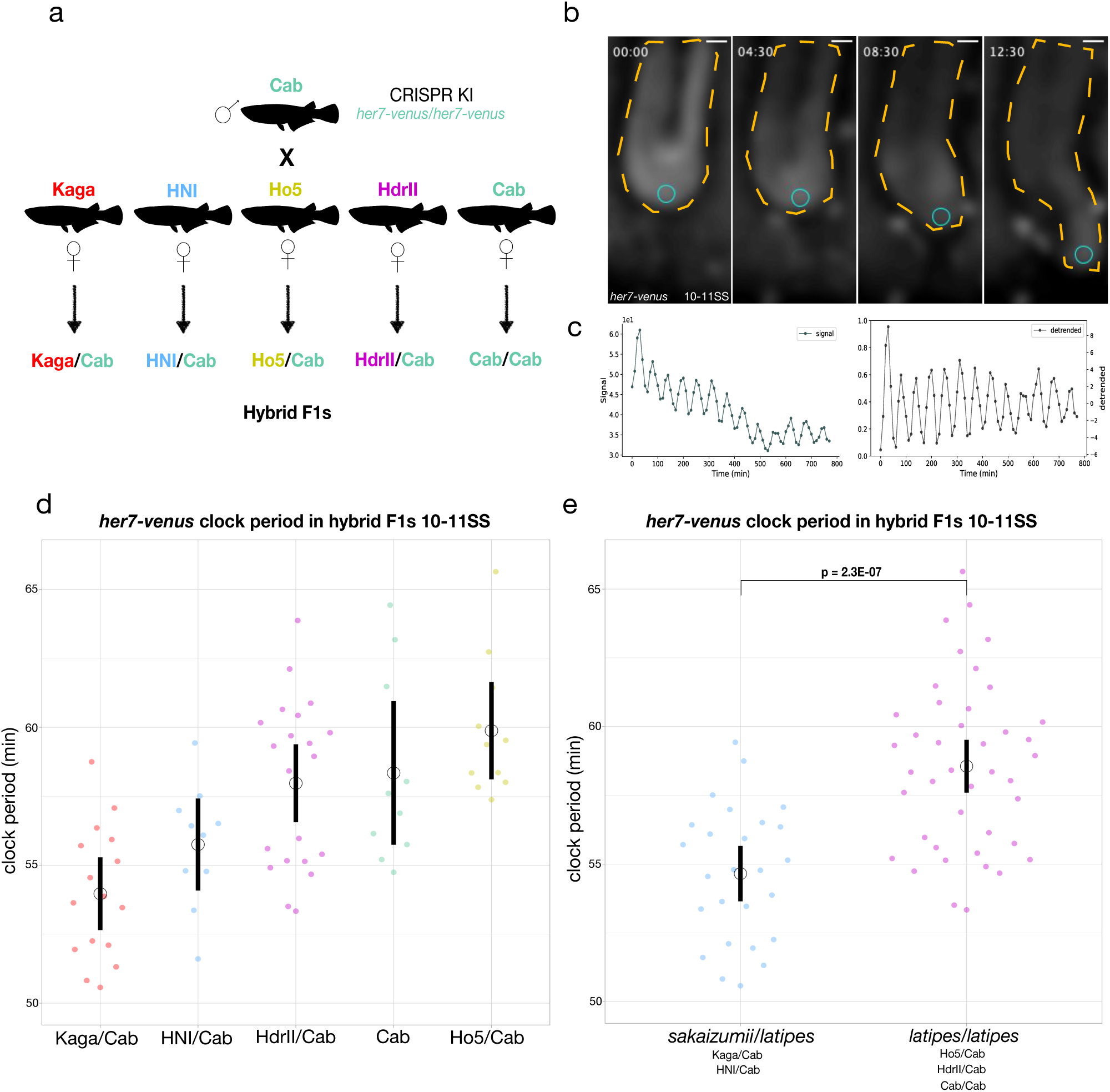
**Hybrid F1 populations show an intermediate clock period timing** (a) Schematic diagram of genetic crosses performed to generate hybrid F1 fish. A male Cab homozygous for the endogenously tagged clock oscillator *her7-venus* is crossed to *wt* Kaga, HNI, Ho5 HdrII and Cab females (b) Selected frames from *in-vivo* tail imaging of endogenous *her7-venus* oscillations at the 10-11 somite stage. Blue circle indicates the location of extracted raw intensity grey values for posterior period analysis over the course of imaging. Yellow dotted lines highlight the outlines of the tail tissue. Time in hours. Scale bar = 30 μm. (c) Raw and detrended signal graphs extracted from the mean intensity grey values of *her7-venus* expression in (b) show oscillatory signal. (d) endogenous *her7-venus* clock period measurements in hybrid F1 embryos at the 10-11SS. Kaga/Cab F1 hybrids have the fastest *her7-venus* clock period (53.97 minutes (sd +/- 2.40)), while HNI/Cab F1s show (55.75 minute (sd +/- 2.21)), hybrid F1 Cab/Cab show (58.34 minutes (sd +/- 3.45)), Cab/HdrII show (57.97 minutes (sd +/- 3.24)), Cab/Ho5 show (59.88 minutes (sd +/- 2.51)). One-way ANOVA p=2.2E-05. Post-Hoc Tukey HSD testing shows significant differences between the following groups Kaga/Cab and Cab/Cab p adjusted = 2.2E-03, Kaga/Cab and HdrII/Cab p adjusted = 4.0E-04, Kaga/Cab and Ho5/Cab p adjusted = 8.9E-06, HNI/Cab and Ho5/Cab p adjusted =9.7E-03. N= 16 Kaga/Cab F1, N= 10 HNI/Cab F1, N= 10 Cab/Cab F1, N= 21 HdrII/Cab F1, N= 11 Ho5/Cab F1. (e) endogenous *her7-venus* clock period measurements in hybrid F1 of *Oryzias sakaizumii/Oryzias latipes* and *Oryzias latipes/Oryzias latipes* embryos at the 10- 11SS. *Oryzias sakaizumii/Oryzias latipes* F1 hybrids have a faster *her7-venus* clock period (54.65 minutes (sd +/- 2.45)) than hybrid *Oryzias latipes/Oryzias latipes* embryos (58.56 minutes (sd +/- 3.05)). Welch two sample t-test p = 2.3E-07. N= 26 *Oryzias sakaizumii/Oryzias latipes* F1 hybrid embryos, N= 42 *Oryzias latipes/Oryzias latipes* F1 hybrid embryos.

## Modular control of segmentation timing and size

We quantified the segmentation clock period in 638 F2 embryos and found a wide distribution of timings that exceeded those occurring in the parental populations (Fig. 3a, Extended Data Fig. 4a-c). The statistical analysis of variances in the parental *vis-a-vis* F2 samples indicated equality (F-test for equality of variances p=0.42 and 0.11) and therefore the wide distribution we report in the F2s is evidence of transgressive segregation^29,30^. The distribution of segmentation clock periods in the F2 offspring was continuous from the fastest (47.1 minutes) to the slowest (69.2 minutes), showing a 22.1 minutes period difference (Fig. 3a, Extended Data Fig. 4d). In addition, we measured PSM and nascent somite size in the F2 embryos, and like segmentation timing, we found a continuous distribution and a wide phenotypic range that exceeded the parental extremes in both directions (Fig. 3b-c, Extended Data Fig. 4e-f). Taken together the data obtained from this F2 cross argues for the polygenic nature and complex genetic control of segmentation timing, PSM size and somite size.

**Figure 3:**
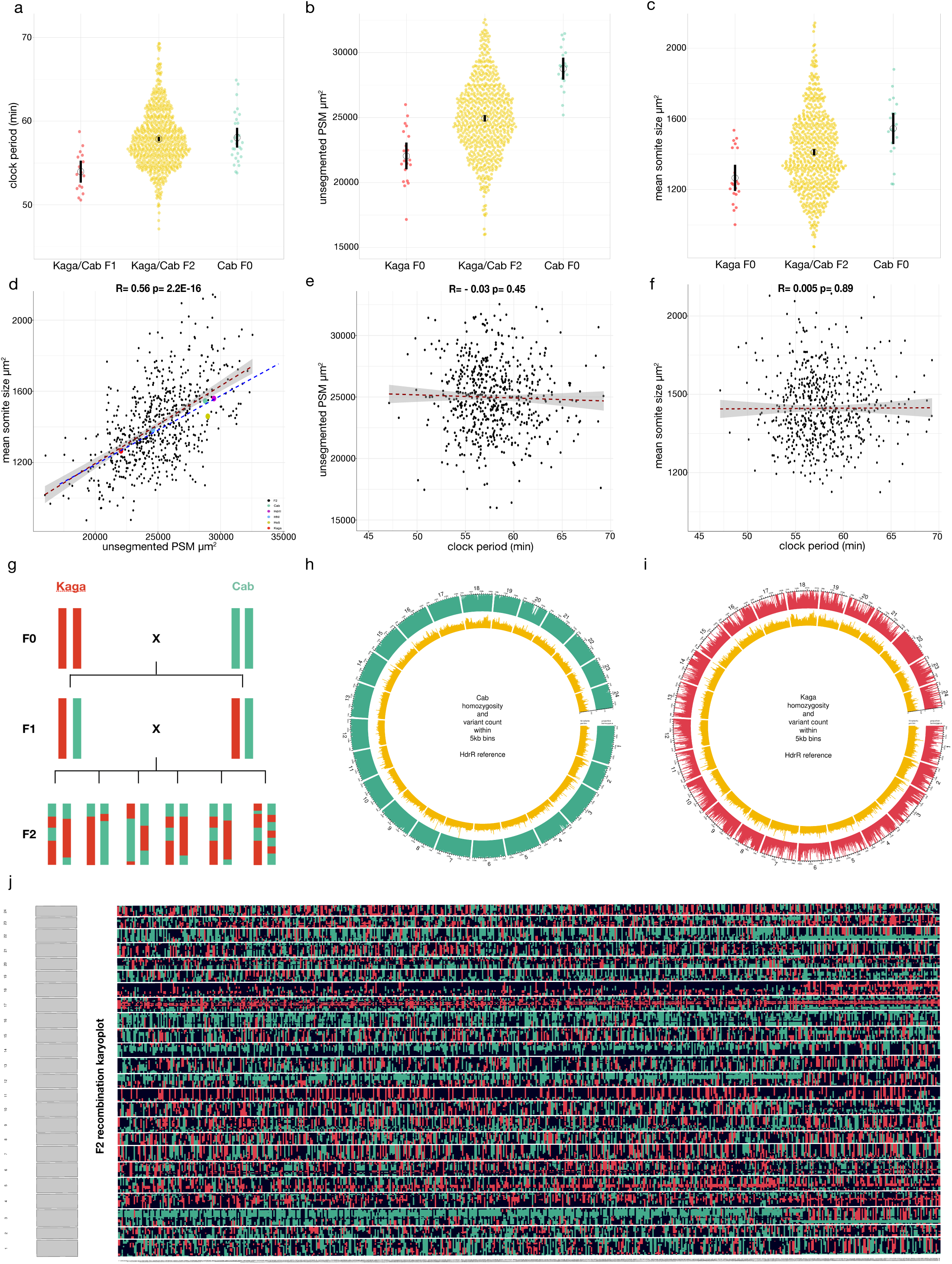
**Genotype-phenotype map of segmentation timing and size in an F2 Kaga/Cab cross** (a) endogenous *her7-venus* clock period measurements in F2 Kaga/Cab embryos (57.86 minutes (sd +/- 3.43)), F1 Kaga/Cab (53.97 minutes (sd +/- 2.4)) and F0 Cab/Cab (58.03 minutes (sd +/- 3.02)). Quantifications were done at the 10-11SS. Black circle = mean period. Black line = 95% confidence interval. Each dot is one embryo. N= 638 F2 Kaga/Cab embryos N= 16 F1 Kaga/Cab embryos N= 28 F0 Cab embryos. (b) unsegmented PSM size in F2 Kaga/Cab (mean = 24944μm^2^ (sd +/- 2930)) as compared to the parental F0 Cab (mean=28770μm^2^ (sd +/- 1709)) and Kaga (mean=22048μm^2^ (sd +/- 2145)). N= 19 Cab, N= 20 Kaga, N= 633 Kaga/Cab F2 (c) mean somite area of nascent somites in Kaga and Cab F0 embryos at the 10-11SS compared to the F2 Kaga/Cab cross. Average sizes: F2 Kaga/Cab (1411μm^2^ (sd +/-222)), Kaga (1265μm^2^ (sd +/- 154)), Cab (1545μm^2^ (sd +/- 177)) . N= 20 Kaga F0 embryos, N= 19 Cab F0 embryos, N= 631 Kaga/Cab F2 embryos. (d) Pearson’s correlation of mean somite size and unsegmented PSM size across all F2 Kaga/Cab embryos R= 0.56 p-value= 2.2E-16. Red dotted line= linear fit for F2 data, grey shaded area= 95% confidence interval for F2 data, slope = 0.044. Blue dotted line= linear fit for F0 average data (Cab, HdrII, HNI, Ho5, Kaga, coloured dots), slope= 0.046 N= 631 Kaga/Cab F2 (black dots) (e) Pearson’s correlation between clock period and unsegmented PSM size in F2 Kaga/Cab cross R= -0.03 p -value= 0.45, N= 623 Kaga/Cab F2. (f) Pearson’s correlation between mean somite size and clock period across all F2 Kaga/Cab embryos R= 0.005 p-value= 0.89. Red dotted line= linear fit, grey shaded area= 95% confidence interval N= 623 Kaga/Cab F2. (g) schematic diagram of genetic crosses to generate F2 Kaga/Cab embryos showing only one chromosome and assuming homozygosity across all sites. Crossing Kaga F0 to Cab F0 generates a hybrid F1 that is heterozygous at all sites. Incrossing F1 Kaga/Cab hybrids generates an F2 population with each individual having a unique genetic composition due to the random nature of meiotic recombination events. (h) circos plot showing the 24 chromosomes of whole genome sequenced F0 Cab embryos aligned against the HdrII reference genome for southern medaka populations. Proportion of homozygous SNPs within 5kb bins in the Cab F0 genome is shown in green and number of SNPs in each bin in yellow. The mean homozygosity across all bins is 83% (i) circos plot showing the 24 chromosomes of a whole genome sequenced F0 Kaga embryo aligned against the HdrII reference genome. Proportion of homozygous SNPs within 5kb bins in the Kaga F0 genome is shown in red and number of SNPs in each bin in yellow. The mean homozygosity across all bins is 31%(j) recombination karyoplot for all 24 chromosomes of whole genome sequenced F2 Kaga/Cab embryos. The plot is based on the ratio of reads mapping to either the Cab or Kaga allele within 5-kb bins (details in Methods). Homozygous Cab blocks are shown in green, heterozygous loci are shown in black and homozygous Kaga loci are shown in red. N = 600 F2 Kaga/Cab embryos.

Interestingly, while further analysing the considerable variation in spatial measures observed in F2 embryos, we found a correlation of nascent somite size to PSM size that mirrored the spatial scaling found across the five different inbred strains (Fig. 3d). The maintenance of a linear relationship between PSM and somite size even in F2 embryos provides evidence for a developmental constraint mechanism^31,32^ underlying segment scaling. Importantly, however, we found that the correlation between PSM/nascent somite size and segmentation clock period, which we had seen across the inbred strains, is absent in the F2 data (Fig. 3e-f, Extended Data Fig. 4g-l). These results indicate that segmentation timing and size can be functionally decoupled.

To link the phenotypes to the underlying genetics we performed *dev*QTL mapping in F2 embryos. This was based on whole genome sequencing (WGS) of the parental Cab and Kaga fish with high coverage (26x and 29x, respectively) (Fig. 3g-i, Extended Data Fig. 5a). We observed a high level of homozygosity (83% of all loci genome-wide) in Cab, while in Kaga, homozygosity was lower (31% homozygosity across all loci genome-wide) (Fig. 3h-i). In total, we identified ∼2.2 million homozygous divergent SNPs between Kaga and Cab that segregated as expected in the F1 Kaga/Cab hybrids (Extended Data Fig. 5b-d). To uncover the genetic basis underlying segmentation timing and size control we built a genome-level genetic recombination map for every F2 embryo by whole genome sequencing (WGS) of 600 F2 embryos with low coverage (∼1x) (Fig. 3j). In conjunction with deep sequencing of the parental populations and hybrid F1s (Fig. 3h-i, Extended Data Fig. 5a-d) we were able to assign one of three genotypes (homozygous Cab, heterozygous Cab/Kaga, homozygous Kaga) to every genomic position for each F2 embryo (Figure 3j, Extended Data Fig. 5e-f, Methods). Combining our quantitative phenotypic measurements with the genomic information of individual F2 embryos we performed *developmental* QTL (*dev*QTL) mapping on both traits.

## *dev*QTL mapping and functional validation on segmentation timing and PSM size

We used the Genome-Wide Complex Trait Analysis (GCTA)^33^ implementation of a linear mixed model to map the *dev*QTLs in F2 embryos associated with segmentation timing (Methods). This revealed several genomic loci that passed the significance threshold, located on chromosomes 3, 4 and 10 (Fig. 4a). These regions contained a total of 46,872 single nucleotide polymorphisms (SNPs) that were homozygous-divergent in the F0 parental strains (Cab and Kaga), the majority of which occurred in non-coding or intergenic portions of the genome (Extended Data Fig. 6a-b). Overall, the SNPs fell within the genomic coordinates of 57 genes. To further refine our search for candidates, we assayed which of the genes are transcriptionally active within the PSM, using bulk-RNA sequencing. From the list of 35 expressed genes and based on GO analysis^34^, we selected a subset of seven candidate genes for *in-vivo* functional validation. We employed an F0 CRISPR/Cas9 knock-out approach^35–38^ and quantified segmentation clock period using *her7-venus in-vivo* imaging (Fig. 4b). Two out of seven targeted genes, i.e. *mesoderm posterior b* (*mespb)* on chromosome 3 and *paraxial protocadherin 10b* (*pcdh10b)* on chromosome 10, showed a minor but significant decrease of segmentation clock period (Fig. 4b, Extended Data Fig. 7a-d). Hybridisation chain reaction (HCR) on both genes in the parental Kaga and Cab strains showed expression domains in the unsegmented PSM (Extended Data Fig 7e-h"). To assess whether the effects of both genes are additive, we performed combinatorial targeting of *mespb/pchd10b* (Fig. 4b). Our results showed a phenocopy of the single knock-outs, suggesting the effects are non-additive and could be mediated through a common genetic pathway.

**Figure 4:**
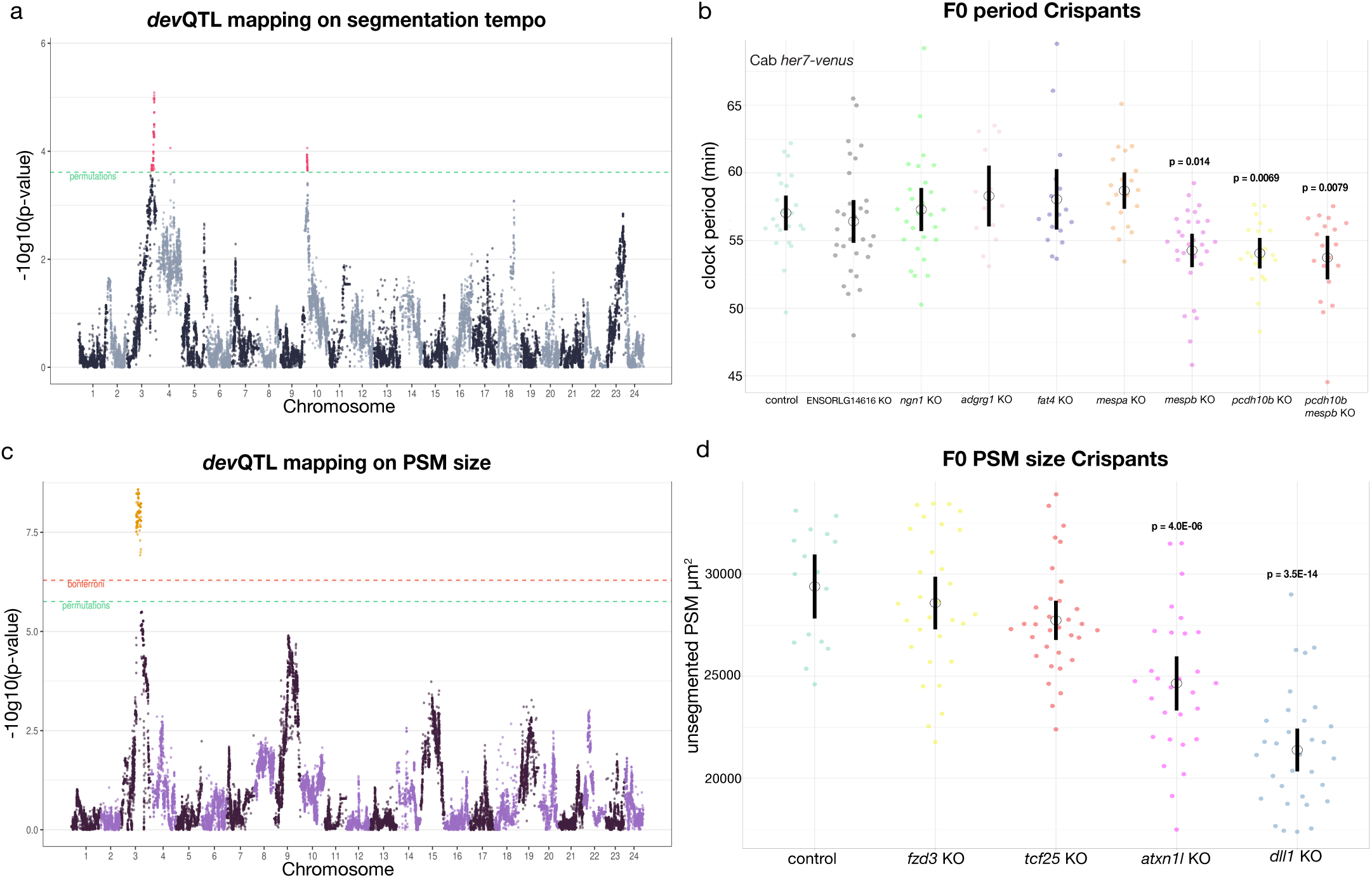
***dev*QTL mapping and functional validation of segmentation timing and PSM size** (a) *dev*QTL on segmentation timing performed on the F2 Kaga/Cab cross shows loci that passed the significance threshold located on chromosomes 3, 4 and 10. Manhattan plot of the genetic linkage results for the inverse-normalised clock period phenotype. Pseudo-SNPs with p-values lower than the permutation significance threshold are highlighted in red. GCTA-LOCO mixed linear model was used, significance threshold was set at 10 permutations (red) (for details see Methods). (b) endogenous *her7-venus* clock period measurements in Control, *ENSORLG14616, ngn1, adgrg1, fat4, mespa, mespb*, *pcdh10b, mespb*+*pcdh10b* F0 Cab Crispants imaged at the 10-11SS. Kruskal-Wallis’ test p= 8.88E-08. Post-Hoc Dunn’s test: only *mespb* (p adjusted=1.4E-02), *pcdh10b* (p adjusted=6.9E-03)*, mespb*+*pcdh10b* (p adjusted=7.9E-03) show a significant difference in clock period as compared to control injected embryos. N= 23 control injected Cab *her7-venus* embryos, N= 30 *ENSORLG14616,* N= 27 *ngn1,* N= 13 *adgrg1,* N= 17 *fat4,* N= 20 *mespa,* N= 29 *mespb,* N= 20 *pcdh10b,* N= 19 *pcdh10b+mespb* CRISPR/Cas9 injected into Cab *her7-venus.* (c) *dev*QTL on unsegmented PSM size performed on the F2 Kaga/Cab cross shows a single locus that passed the significance threshold located on chromosome 3 at a distinct genomic coordinate from that found on chromosome 3 for the segmentation timing *dev*QTL hits. Manhattan plot of the genetic linkage results for the unsegmented PSM size phenotype. Pseudo-SNPs with p-values lower than the permutation (green) and Bonferroni (red) significance thresholds are highlighted in orange. GCTA-LOCO mixed linear model was used, significance threshold was set at 10 permutations and Bonferroni p-value was set by dividing 0.05 by the number of pseudo-SNPs in the model (for details see Methods). (d) unsegmented PSM size Cab F0 CRISPR/Cas9 knock-outs performed on candidate genes from the *dev*QTL mapping results in (c) on Control, *fzd3*, *tcf25, atxn1l* and *dll1* F0 Cab Crispants imaged at the 10-11SS. Both *atxn1l* and *dll1* Crispants showed significantly smaller unsegmented PSM size (24645 µm^2^ (sd +/- 3426)) and (21382 µm^2^ (sd +/- 2868)) respectively than Cab control Cas9 mRNA injected embryos (29392µm^2^ (sd +/- 2851)). One-way ANOVA test p= 7.5E-15. Post-Hoc Dunnett’s test: only *atxn1l* (p=4.0E-06), *dll1* (p=3.5E-14) show a significant difference in PSM size as compared to control injected embryos. N= 16 control injected embryos, N = 30 *fzd3* Crispants N= 33 *tcf25* Crispants N= 29 *atxn1l* Crispants N= 32 *dll1* Crispants.

The *dev*QTL mapping using PSM size as a linked trait identified a single significant region on chromosome 3 (Fig. 4c). This region is distinct from the one we obtained on the same chromosome for segmentation timing. In this region, we identified 204 genes, of which 155 are expressed in the PSM. As a proof of principle, we selected 4 candidate genes based on GO annotation and performed CRISPR/Cas9 loss-of function analysis in F0 embryos (Extended Data Fig. 8e-h). We were able to identify two Crispants, i.e. *atxn1l* (a notch co-factor) and *dll1* (a notch ligand), that showed a reduction in PSM size compared to control embryos (Fig. 4d, Extended Data Fig. 8a-h). In both *atxn1l* and *dll1* Crispants we assayed whether there was an effect on segmentation timing. We were able to extract reliable segmentation period measurements only from a subset of *atxn1l* and *dll1* Crispant embryos, likely due to an overall down regulation of *her7* expression, however, the results showed that segmentation period does not differ from control embryos for either Crispant (Extended Data Fig. 8c-h), despite a reduction of PSM size. The CRISPR/Cas9 knock-out approach provides proof of principle validation that the *dev*QTL mapping identified functionally relevant genomic regions linked to the control of segmentation timing and PSM size, respectively.

In this study we were able to reveal, using genetic crosses of closely related medaka species, evidence for a developmental constraint mechanism underlying segment size scaling, forming a single functional module. Moreover, we discovered the modular nature of temporal and spatial control of axis segmentation. Interestingly, we found evidence these modules are, effectively, coupled in the natural context, which is reflected by the linear correlation of spatial and temporal developmental traits that we found in the interspecies comparison. This points towards the existence of selective constraints that couple these modules, thereby restricting the phenotypic outcomes realised in the natural setting. Future investigations will aim to understand how selective pressures and developmental modules are integrated, taking into account the environmental context and life-history trade-offs.

## Methods

### Animal husbandry and ethics

Medaka *Oryzias latipes* (Cab, HdrII, Ho5), *Oryzias sakaizumii* (HNI and Kaga) strains^39–41^ and *her7-venus* Cab^5^, were maintained as closed stocks in a constant recirculating system at 27-28°C, with a 14hr light / 10hr dark cycle in the EMBL Laboratory Animal Resources (LAR) fish facility. Animal experiments were performed after project approval by the EMBL Institutional Animal Care and Use Committee (IACUC), IACUC project code is 20/001_HD_AA.

### Live-imaging sample preparation

Embryos were prepared for live-imaging as previously described^42,43^. 1X Tricaine (Sigma-Aldrich #A5040-25G) was used to anaesthetise dechorionated medaka embryos (20 mg/ml - 20X stock solution diluted in 1XERM). Anaesthetised embryos were then mounted using low melting agarose (0.6 to 1%) (Biozyme Plaque Agarose #840101). Imaging was done in 8-well glass-bottomed dishes (Lab-Tek Chambered #1 Borosilicate Coverglass System 155411, Thermo Scientific Nunc, USA).

### Tail explants

Dechorionated Cab and Kaga embryos at the 15-16 somite stage were placed in Gibco CO_2_ independent medium (Thermo Fisher #18045054). Tails were cut 4-5 somites directly above the presomitic mesoderm (PSM). Dissected tails were then placed in individual wells of 8-well glass-bottomed dishes (Lab-Tek Chambered #1 Borosilicate Coverglass System 155411, Thermo Scientific Nunc, USA) with 200µl of Gibco CO_2_ independent medium (Thermo Fisher #18045054).

### Hybridization chain reaction (HCR)

Cab and Kaga embryos at the 10 somite stage were fixed in 4% PFA in PtW for 12-24 hours at 4°C. Embryos were then washed 3x in PtW followed by dehydration and storage in MeOH at -20°C. Medaka *mespb, pcdh10b* and *dll1* probes for hybridization chain reaction^44^ were ordered from Molecular instruments (MI) and the protocol was carried out according to the manufacturer’s guidelines. HCR amplifiers used were B1-546, B2-546, B2-647. Hoechst 33342 (Thermo Fischer #H3570) was used with a dilution of 1:500 of 10 mg/ml stock solution as a nuclear label.

### Microscopy

All embryo screening was done on a Nikon SMZ18 fluorescence stereoscope equipped with a camera. ALC and HCR image acquisition was done either on Nikon SMZ18 fluorescence stereoscope equipped with a camera or on a laser-scanning confocal Leica SP8 (CSU, White Laser) microscope, 20x and 40x objectives were used during image acquisition depending on the experimental sample. For the SP8 confocal equipped with a white laser, the laser emission was matched to the spectral properties of the fluorescent protein of interest. For all F0, F1, F2 imaging of embryos and for all CRISPR/Cas9 knock-outs (KOs) image acquisition was performed using two Zeiss LSM780 laser-scanning confocal microscopes with a temperature control box and an Argon laser at 488 nm, imaged through a 20x plan apo objective (numerical aperture 0.8). All embryos were imaged at 10-11SS unless otherwise stated. Temperature on the incubator box of both microscopes was set at 30°C. To account for slight differences in actual temperature in the wells between the two microscopes used for the F2 data acquisition: mean and intercept period measurements were normalised and plotted by microscope, normalisation was done either by using inverse normalisation with the following formula:

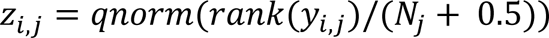

Where *rank*(*y_i_*_,*j*_) is the sample rank of observation *i* within microscope *j*, *N_j_* is the sample size for microscope *j*, and *qnorm* calculates the percentile value based on the normal distribution^45^ or by equating the mean period of all samples on one microscope (reference) to the mean period of all samples on the other microscope. The difference between the mean measurements on the two microscopes translates to 3.5 minutes for mean period and 4.0 minutes for intercept period. For all other experiments either one microscope (reference) was used or a temperature sensor probe GMH 3200 series Thermocouple (Greisinger) was placed into the imaging wells and temperature was measured throughout imaging to ensure equivalent temperatures were measured between the two microscopes.

### Data analysis

Open-source ImageJ/Fiji software^46^ was used for analysis and editing of all images post image acquisition. Stitching was performed using 2D and 3D stitching plug-ins on ImageJ/Fiji. For extracting quantitative values of posterior period oscillations in the endogenous *her7-Venus* line, the time-series movie was first gaussian blurred (sigma 8) in ImageJ/Fiji then ROI manager was used to define fluorescence intensity within a circle (area 300-600μm^2^), at the posterior most tip of the tail, the circle was manually moved to track the movement and growth of the tails over the course of imaging, fluorescent intensity measurements were then concatenated for every time point and extracted from the time-series using a custom made Fiji macro-script provided in Supplementary Information S1. For F0 segmentation time estimation in the Cab, Kaga, HNI, Ho5 and HdrII strains we used segment boundary formation obtained from bright-field live-imaging to determine segment forming time. The time it took to form 5 consecutive segments was calculated for each embryo to get an estimation of segmentation time. For presomitic mesoderm (PSM) size area measurements were done using polygon selection in Fiji on brightfield tail images of 10-11SS embryos on the unsegmented tissue at the tip of the tail. Presomitic mesoderm (PSM) length measurements were done using the segmented line tool in Fiji on brightfield tail images of 10-11SS embryos on the unsegmented tissue at the tip of the tail. Somite length and area measurements were done on the first pair of nascent somites using the line or polygon tool in Fiji on brightfield tail time-lapse imaging of 10-11SS embryos. For volumetric egg measurements the eggs were approximated as oblate spheroids^41^ and the following equation was used *V=4/3.7r.(b)*^2^*.c.* where *V*= volume *b*= semi-major axis and *c*= semi-minor axis. Data was plotted using *ggplot2* and *gganimate* in R software or using *PlotTwist*^47^ and *PlotsofData*.^48^ Pearson’s product moment correlations, F-test for equality of variances, Welch two sample t-tests, Kruskal-Wallis test, one-way ANOVA test, Dunnet’s test, Tukey HSD test and Dunn’s test were all calculated and plotted in R version 4.2.2.

### Wavelet analysis and period extraction

Raw fluorescent intensity measurements were used for wavelet analysis and period extraction using PyBoat^49^. The following settings were used for all samples: sampling interval 10 minutes, cut-off period was set at 100 minutes, window size was set at 150 minutes, the smallest period was set at 40 minutes, the number of periods to scan was set at 200, the highest period was set at 100 mins, detrended signal and normalisation with envelope was used on all samples. The continuous maximum ridge connecting the wavelet power was then plotted. Data within the COI (cone of influence) were then extracted to get period, phase, amplitude and power for each analysed sample. Period values, one for each 10 minute sampling interval, for a total of 300 minutes were then used for all subsequent analysis, either to generate mean period plots (average period in 300 minute interval) or intercept period plots (y-intercept of fitted line on 300 minute interval period measurements). The clock period shown in the main figures corresponds to intercept period measurements unless otherwise stated. Both mean and intercept period measurements are shown in the Extended Data.

### Bulk RNA-sequencing

Dechorionated Cab and Kaga embryos at the 12-13 somite stage were placed in Gibco CO_2_ independent medium (Thermo Fisher #18045054). Using a forceps and a scalpel, tails were cut directly at the unsegmented presomitic mesoderm (PSM) border. Five dissected tails in three replicates for Cab embryos and five dissected tails in three replicates for Kaga embryos were then disrupted and homogenised for total RNA extraction using RNeasy Plus Micro Kit (Qiagen #74034) following the manufacturer’s guidelines. The integrity and concentration of the extracted RNA was checked by using Agilent Bioanalyzer with the RNA 6000 Nano Assay kit. Libraries were prepared using the NEBNext Ultra II Directional RNA Library Prep Kit for Illumina (New England Biolabs) together with the NEBNext Poly(A) mRNA Magnetic Isolation Module (New England Biolabs) using the long inserts version of the manufacturer’s protocol. Briefly, these modifications consisted of 7 minutes of fragmentation, 50 minutes extension for the first strand synthesis, 1:100 dilution of the adaptor and size selection for a 400 base pair long insert. The libraries were quantified using the Qubit HS DNA assay as per the manufacturer’s protocol. For the measurement, 1ul of sample in 199ul of Qubit working solution was used. The quality and molarity of the libraries was assessed using Agilent Bioanalyzer with the DNA HS Assay kit as per the manufacturer’s protocol. The assessed molarity was used to equimolar combine the individual libraries into one pool for sequencing. The pool was sequenced on Illumina NextSeq2000 (Illumina, San Diego, CA, USA) using a P3 flowcell and reading 2 x 150 bases. Sequencing files were demultiplexed using FastQC (version 0.11.9) and the output was collated using MultiQC (version1.10)^50^. Sequencing reads were aligned using STAR (version 2.7.9a)^51^ to the medaka genome (version ASM223467v1) with default parameters. The gene count tables were computed during the alignment with STAR on the medaka gene model (version ASM223467v1.103). Gene counts table is provided as Supplementary Information SI2.

### CRISPR/Cas9 Knock-outs

Embryos (WT Cab or *her7-venus* Cab) were injected at the 1 cell stage. Cas9 mRNA was obtained from pCS2-Cas9 (Addgene #47322) as previously described^52,53^. In-vitro transcription was carried out using mMachine SP6 Transcription Kit (Invitrogen #AM1340) following the manufacturer’s guidelines. RNA cleanup was carried out using RNAeasy Mini Kit (Qiagen #74104). 2-3 synthetic gRNAs per gene targeting exonic regions were designed using CCTop^54^ and ordered from Sigma-Aldrich (spyCas9 sgRNA, 3 nmol, HPLC purification, no modification). A list of all gRNAs and their corresponding target genes used in this study is provided in Supplementary Information SI3. The injection mix consists of 15 ng/µl for each gRNA and 75 ng/µl Cas9 mRNA. For control injections only the Cas9 mRNA was injected.

### DNA extraction and library preparation for F0 Cab and Kaga and F1 Kaga/Cab hybrid samples

DNA from two separate stage 42 embryos of F0 Cab and F0 Kaga and one stage 42 embryo of F1 hybrid Kaga/Cab was extracted using DNeasy Blood and Tissue Kit (Qiagen #69504) following the manufacturer’s guidelines. The extracted DNA was quantified using the Invitrogen Qubit Flex, with the Qbit dsDNA High Sensitivity assay as per the manufacturer’s protocol. For the measurement, 1ul of sample in 199ul of Qubit working solution was used. Samples were then standardised and 1 µg of material was used as an input for library preparation of both F0 Cab and F0 Kaga, while the input amount was 750 ng for the F1 Kaga/Cab hybrid sample. Libraries were prepared using the NEBNext UltraII DNA Library Prep Kit (New England Bioalbs) with NEBNext Multiplex Oligos for Illumina (Unique Dual Index UMI Adaptors DNA Set 1) according to the manufacturer’s instructions without PCR. A size selection for 500 base pairs insert size was performed. The libraries were quantified using the Qubit HS DNA assay as per the manufacturer’s protocol. For the measurement, 1ul of sample in 199ul of Qubit working solution was used. The quality and molarity of the libraries was assessed using Agilent Bioanalyzer with the DNA HS Assay kit as per the manufacturer’s protocol. The assessed molarity was used to equimolar combine the individual libraries into one pool for sequencing for the F0 Cab and F0 Kaga samples. The pool (F0 samples) and the hybrid F1 sample were both sequenced on Illumina NextSeq500 (Illumina, San Diego, CA, USA) using a MID output kit and reading 2 x 150 bases. The sequencing data for this study have been deposited in the European Nucleotide Archive (ENA) at EMBL-EBI under accession number PRJEB59222 (https://www.ebi.ac.uk/ena/browser/view/PRJEB59222).

### DNA extraction and library preparation for F2 samples

After imaging, F2 embryos were recovered and grown in individual wells of 12- well plates (Thermo-Fisher #150628) until stage 41-42. Samples were then placed in 1.5ml Eppendorf tubes, snap-frozen and stored in -80°C. For DNA extraction of all F2 samples, the following protocol was used: 40µl Finclip Buffer was added to snap-frozen embryos. Embryos were then Incubated at 65 °C overnight. 80µl distilled H_2_O was then added and samples were incubated for 10 minutes at 95 °C. Samples were then centrifuged at 10,000 rpm at 4°C for 25 minutes. Supernatant containing genomic DNA was then used for subsequent library preparation. Fin clip buffer is composed of 100 ml 2M Tris pH 8.0, 5 ml 0.5M EDTA pH 8.0, 15 ml 5M NaCl, 2.5 ml 20% SDS, H_2_O to 500 ml, sterile filtered. The extracted DNA per sample was quantified using the Invitrogen Qubit Flex, with the Qbit dsDNA High Sensitivity assay as per the manufacturer’s protocol. For the measurement, 1ul of sample in 199ul of Qubit working solution was used. Each sample was then standardised with water to a final concentration of 2ng/ uL. Sequencing libraries were prepared as previously described ^55,56^ using the automated liquid handler Biomek i7 system (Beackman Coulter). In short, 1.25 uL of each sample was taken into a tagmentation reaction containing 1.25 uL of Dimethylformamide, 1.25 uL of tagmentation buffer (40 mM Tris-HCl pH 7.5, 40 mM MgCl_2_) and 1.25 uL of an in-house generated and purified Tn5^56^ and diluted to a 1:100 ratio with water. The mixture was incubated at 55°C for 3 minutes. After that, 1.25 uL of SDS 0.2% was added to stop the tagmentation reaction followed by a 5 minutes incubation at room temperature. The resulting fragments were then amplified by PCR using 6.75 uL of KAPA 2 X HiFi master mix, 0.75 uL of Dimethyl sulfoxide and 2.5 uL of dual indexed primers. Cycling conditions were as follows: 3 minutes at 72°C; 30 seconds at 95°C; 12 cycles of 20 seconds at 98°C, 15 seconds at 58°C, 30 seconds at 72°C; 3 minutes at 72°C. The resulting PCR products were then pooled together by combining 4uL of each desired sample. Finally, the pools were size selected using two rounds of magnetic SPRI bead purification (0.6X) and quantified using the Qbit dsDNA High Sensitivity assay. Samples were sequenced either on Illumina Hiseq4000 (150 paired end) (Illumina, San Diego, CA, USA) or Illumina Nextseq2000 (150 paired end) (Illumina, San Diego, CA, USA). All sequencing data is deposited at the ENA under study number:

### Whole genome sequencing, alignment, and variant calling for Cab and Kaga F0 and Kaga/Cab hybrid F1s

Coverage for each sample was measured using SAMtools^57^ with a mean of ∼26x for F0 Cab, ∼29x for F0 Kaga and ∼59x for Kaga/Cab hybrid .Reads were then aligned to the medaka HdrR reference (Ensembl release 104, build ASM223467v1) using *BWA-MEM2*,^58^ sorted the aligned .sam files, marked duplicate reads, merged the paired reads with the *Picard* toolkit,^59^ and indexed the .bam files with *SAMtools*.^60^ The Snakemake pipeline used to map and align these samples can be found in the GitHub repository here: https://github.com/birneylab/somites. To call variants, we followed the *GATK* best practices (to the extent they were applicable)^61–63^ with *GATK*’s HaplotypeCaller and GenotypeGVCFs tools,^64^ then merged all calls into a single .vcf file with *Picard*^59^ Finally, we extracted the biallelic calls for Cab and Kaga with bcftools,^57^ counted the number of SNPs within non-overlapping, 5-kb bins, and calculated the proportion of SNPs within each bin that were homozygous to generate Fig. 3h-i and Extended Data Fig. 7 using R version 4.2.2,^65^ the *tidyverse* suite of R packages,^66^ and *circlize*.^67^ To assess whether the low homozygosity observed in the Kaga strain was caused by a reference bias, we also aligned the reads of the Kaga F0 sample to the northern Japanese HNI reference (Ensembl release 105 build ASM223471v1) to generate Extended Data Fig. 5c using the same process as above.

### Whole genome sequencing and alignment on Kaga/Cab F2s

For all F2 samples the sequencing reads were aligned to the HdrR reference (Ensembl release 104, build ASM223467v1) using *BWA-MEM2*,^58^ sorted the aligned .sam files, marked duplicate reads, merged the paired reads with the *Picard* toolkit^59^ and indexed the .bam files with *SAMtools*^60^ in the same manner as for the F0 and F1 samples. To map the aligned F2 sequences to the genomes of their parental strains, we selected only biallelic SNPs that were homozygous-divergent in the F0 generation (that is, homozygous for the reference allele in Cab and homozygous for the alternative allele in Kaga, or vice versa) *and* heterozygous in the F1 generation. The number of SNPs that met these criteria per chromosome are set out in Extended Data Fig. 7. To call variants in the shallow-sequenced F2 samples, we applied a method involving the use of a Hidden Markov Model (HMM) to classify regions of the genome as one of the three genotypes (AA, AB, or BB), based on the proportion of reads that supported either parental allele, as described previously.^68^ We accordingly used *bam-readcount*^69^ to count the number of reads that supported either the Cab or the Kaga allele for all SNPs that met the criteria described above (i.e. biallelic SNPs that were homozygous-divergent in the parental Cab and Kaga F0 samples), summed the read counts within 5 kb blocks across the genome, and calculated the frequency of reads within each bin that supported the Kaga allele. This generated a value for each bin between 0 and 1, where 0 signified that all reads within that bin supported the Cab allele, and 1 signified that all reads within that bin supported the Kaga allele. Bins containing no reads were assigned a value of 0.5. We then used these values for all F2 individuals as the input to an HMM with the software package *hmmlearn*,^70^ which we applied to classify each bin as one of three states, with state 0 corresponding to homozygous-Cab, state 1 corresponding to heterozygous, and state 2 corresponding to homozygous-Kaga. Across each chromosome of every sample, the output of the HMM was expected to produce a sequence of states. Based on previous biological knowledge that crossover events occur on average less than once per chromosome^71^, we expected to observe the same state persisting for long stretches of the chromosome, only changing to another state between 0 and 3 times, and rarely more. To achieve this, we adjusted the HMM’s transition probabilities to be nearly 0, and the Gaussian emission probabilities for each state to have a variance of 0.8, which resulted in long "blocks" of the same genotype call across the chromosome with only a small number of average transitions (i.e. crossover events) per chromosome. We then used the R package *karyoploteR*^72^ to generate the recombination block plot shown in Fig. 3J. Extended Data Fig. 7 shows the proportion of 5-kb bins called as either homozygous-Cab, heterozygous, or homozygous-Kaga within each F2 sample (points). The ordinary expectation for the ratios would be 0.25, 0.5, and 0.25 respectively. However, we observe a skew towards homozygous-Cab and away from homozygous Kaga. This was likely caused by the lower level of homozygosity in Kaga, and also potentially a degree of hybrid incompatibility between Cab and Kaga, given the two strains were derived from populations that are at or beyond the point of speciation. In the downstream analysis, we excluded the 22 samples that showed poor coverage across the genome, leaving *N* = 600 for the genetic association testing. For these remaining samples, we "filled" the bins with missing genotypes based on the call of the previous called bin, or if unavailable (e.g. the missing bin was at the start of the chromosome), then the next called bin. We used these filled genotype calls for the genetic association tests described below.

### *dev*QTL and VEP output

We used the recombination blocks called by the HMM as pseudo-SNPs in an F2- cross *dev*QTL. To detect associations between the pseudo-SNPs and the two phenotypes of interest, we used a linear mixed model (LMM) as implemented in GCTA^33^. For the genetic relationship matrix (GRM), we additionally used the leave-one-chromosome-out implementation of GCTA’s LMM, which excludes the chromosome on which the candidate SNP is located when calculating the GRM. A GRM constructed from the entire genome is presented as a heatmap in Extended Data Fig. 5e with each sample represented on each axis, and lighter colours representing a higher degree of relatedness between a pair of samples. The square in the top right-hand corner is created by samples ∼550-648, which have distinct genotypes to the rest of the samples due to their having been bred from different F1 parents. To set the significance threshold, we permuted the phenotype across samples using 10 different random seeds, together with all covariates when included, and ran a separate linkage model for each permutation. We then set the lowest *p*-value from all 10 permutations as the significance threshold for the non-permuted model. We additionally applied a Bonferroni correction to our *p*-values by dividing *α* (0.05) by the number of pseudo-SNPs in the model, and set this as a secondary threshold. For the pseudo-SNPs (5-kb regions) which returned a lower *p*-value than the significance level set by the permutations, we ran Ensembl’s Variant Effect Predictor to identify the variants that are predicted to be more likely to disrupt the functions of the sequence.

## Data availability

All data are available within the article, supplementary files and source data files. All materials used in this study are available by request from the corresponding author. Source data are provided with this paper. The sequencing data for this study have been deposited in the European Nucleotide Archive (ENA) at EMBL-EBI under accession number PRJEB59222 (https://www.ebi.ac.uk/ena/browser/view/PRJEB59222).

## Code availability

All code used in this study is available at https://github.com/birneylab/somites. The custom-made Fiji macro-script for concatenating and extracting fluorescent intensity measurements from a defined ROI is available in Supplementary Information S1.

## Acknowledgments

We would like to thank all members of the Aulehla lab for discussions throughout the project and Jordi Van Gestel for comments on the manuscript. The European Molecular Biology Laboratory (EMBL-Heidelberg) Genecore is acknowledged for support in WGS data acquisition and analysis. We would like to thank Vladimir Benes and members of his team at Genecore EMBL Heidelberg for continuous help and support, Tobias Rausch for computational work on the WGS data and Mireia Osuna Lopez for help in library preparation of all WGS DNA and RNA-seq data and Jonathan Landry for help with the RNA-seq data analysis. The EMBL-EBI’s high performance computing cluster is acknowledged for providing the computational resources required for the *dev*QTL analysis. We would like to thank Sabine Goergens and her team at EMBL Laboratory Animal Resource (LAR) Team fish facility for their excellent, daily animal husbandry and expert support. We would like to thank the National Bioresource Project (NBRP) Japan for access to medaka hatching enzyme. This work was supported by the European Molecular Biology Laboratory (EMBL), the EMBL interdisciplinary Postdoc (EIPOD4) under Marie Sklodowska-Curie Actions Cofund (grant agreement n.847543) fellowship for funding to A.S., and the EMBL International PhD Programme fellowship to I.B. This work also received support from the European Research Council under an ERC consolidator grant agreement n.866537 to A.A.

## Author Contributions

A.S: Conceptualization, Methodology, Validation, Formal analysis, Investigation, Data Curation, Writing - original draft, Visualisation. I.B: Formal analysis, Methodology, Data Curation, Software, Investigation, Visualization, Writing-review and editing. T.F. Methodology, Formal analysis, Software, Data Curation. C.V. Methodology, Resources, Validation, Writing - review and editing. J.W. Conceptualization, Resources, Writing - review and editing. F.L. Conceptualization, Resources, Writing - review and editing. K.N. Conceptualization, Resources. E.B. Conceptualization, Methodology, Resources, Writing - review and editing, Supervision, Project administration, Funding acquisition. A.A: Conceptualization, Methodology, Resources, Writing - original draft preparation, Supervision, Project administration, Funding acquisition.

## Competing interests

The authors declare that they have no conflict of interest.

Supplementary Information is available for this paper.

Correspondence and requests for materials should be addressed to

## Extended Data legends

**Extended Data Figure 1:**
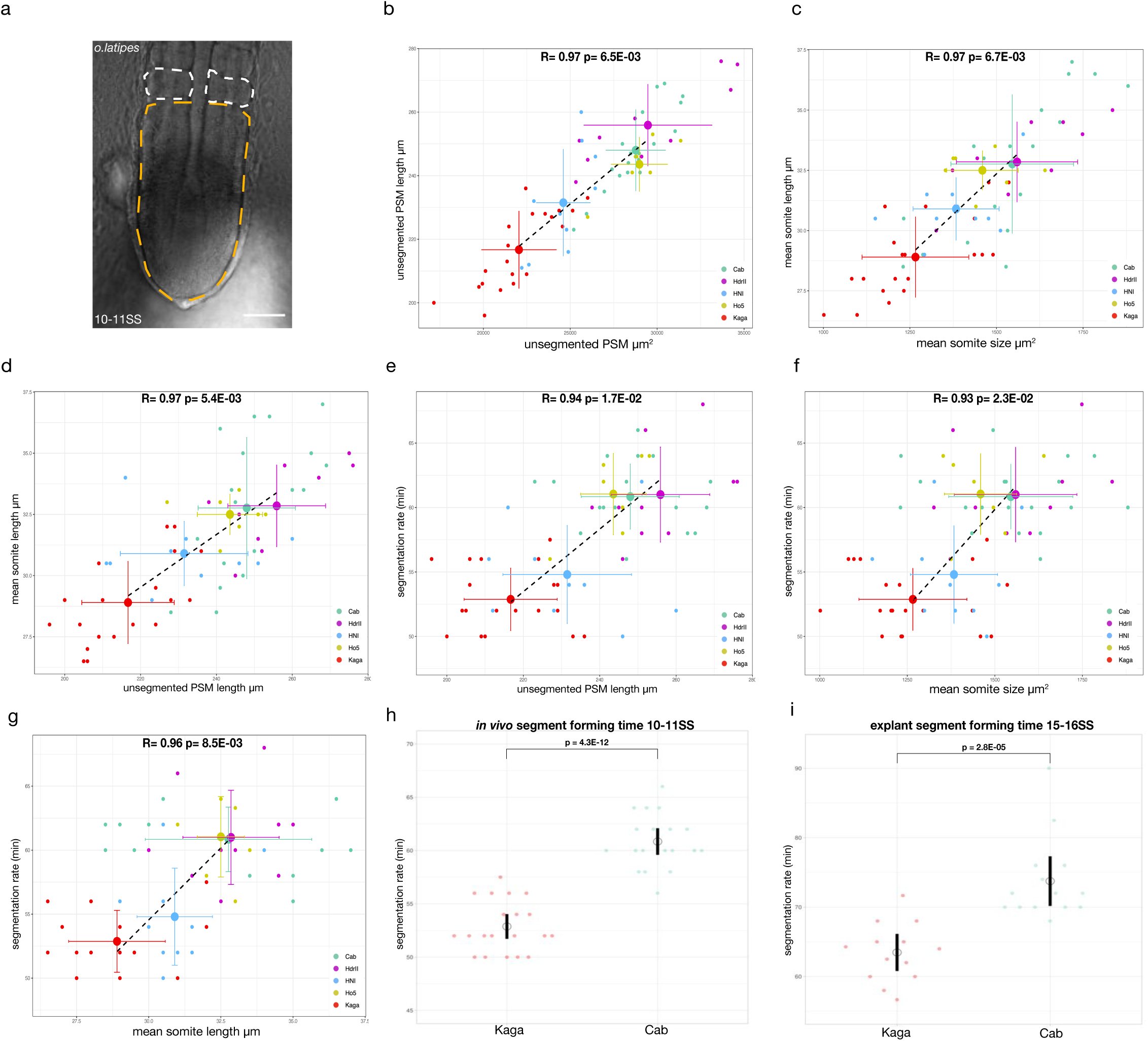
**Scaling of embryonic segmentation timing and PSM/somite size in *Oryzias sakaizumii* and *latipes*** (a) bright-field image of unsegmented presomitic mesoderm (PSM) and nascent somites at the 10-11 somite stage (SS) Cab F0 medaka embryo. Yellow dotted line delineates the unsegmented PSM area while white dotted lines delineate the area of nascent somites. Scale bar= 50µm.(b) Pearson’s correlation of average trait values for unsegmented PSM length and unsegmented PSM area measured at 10-11 somite stage in *Oryzias sakaizumii* Kaga (red), HNI (blue) and *Oryzias latipes* Cab (turquoise), HdrII (magenta), Ho5 (yellow) embryos R= 0.97 p-value= 6.5E-03. Black dotted line= linear fit on average trait values, N= 20 Kaga, N= 10 HNI, N= 19 Cab, N= 10 HdrII, N= 7 Ho5. (c) Pearson’s correlation of average trait values for mean somite size and length measured at 10-11 somite stage in *Oryzias sakaizumii* Kaga (red), HNI (blue) and *Oryzias latipes* Cab (turquoise), HdrII (magenta), Ho5 (yellow) embryos R= 0.97 p-value= 6.7E-03. Black dotted line= linear fit on average trait values, N= 20 Kaga, N= 10 HNI, N= 19 Cab, N= 10 HdrII, N= 7 Ho5.(d) Pearson’s correlation of average trait values for unsegmented PSM length and somite length measured at 10-11 somite stage in *Oryzias sakaizumii* Kaga (red), HNI (blue) and *Oryzias latipes* Cab (turquoise), HdrII (magenta), Ho5 (yellow) embryos R= 0.97 p-value= 5.4E-03. Black dotted line= linear fit on average trait values, N= 20 Kaga, N= 10 HNI, N= 19 Cab, N= 10 HdrII, N= 7 Ho5. (e)Pearson’s correlation of average trait values for unsegmented PSM length and segmentation rate measured at 10-11 somite stage in *Oryzias sakaizumii* Kaga (red), HNI (blue) and *Oryzias latipes* Cab (turquoise), HdrII (magenta), Ho5 (yellow) embryos R= 0.94 p-value= 1.7E-02. Black dotted line= linear fit on average trait values, N= 20 Kaga, N= 10 HNI, N= 19 Cab, N= 10 HdrII, N= 7 Ho5. (f)Pearson’s correlation of average trait values for mean somite size and segmentation rate measured at 10-11 somite stage in *Oryzias sakaizumii* Kaga (red), HNI (blue) and *Oryzias latipes* Cab (turquoise), HdrII (magenta), Ho5 (yellow) embryos R= 0.93 p-value= 2.3E-02. Black dotted line= linear fit on average trait values, N= 20 Kaga, N= 10 HNI, N= 19 Cab, N= 10 HdrII, N= 7 Ho5. (g)Pearson’s correlation of average trait values for mean somite length and segmentation rate measured at 10-11 somite stage in *Oryzias sakaizumii* Kaga (red), HNI (blue) and *Oryzias latipes* Cab (turquoise), HdrII (magenta), Ho5 (yellow) embryos R= 0.96 p-value= 8.5E-03. Black dotted line= linear fit on average trait values, N= 20 Kaga, N= 10 HNI, N= 19 Cab, N= 10 HdrII, N= 7 Ho5. (h) axis segmentation rate from brightfield time-lapse imaging of Kaga and Cab F0 embryos measured at the 10-11 somite stage calculated from the time it takes to form 5 consecutive pairs of somites. Kaga embryos have a faster segmentation period (52.88 minutes (sd +/- 2.42)) than Cab (60.84 minutes (sd +/- 2.52)). Welch two sample t-test p = 4.3E-12. Black circle = mean Black line = 95% confidence interval. N= 20 Kaga embryos, N= 19 Cab embryos. (i) *ex-vivo* explant axis segmentation rate in bright-field imaging of Kaga and Cab F0 embryos at the 15-16 somite stage calculated from the time it takes to form 5-6 consecutive pairs of somites. Kaga embryos have a faster *ex-vivo* segmentation period (63.47 minutes (sd +/- 4.2)) than Cab (73.75 minutes (sd +/-5.9)). Welch two sample t-test p = 2.8E-05. Black circle = mean Black line = 95% confidence interval. N= 13 Kaga embryos, N= 14 Cab embryos.

**Extended Data Figure 2:**
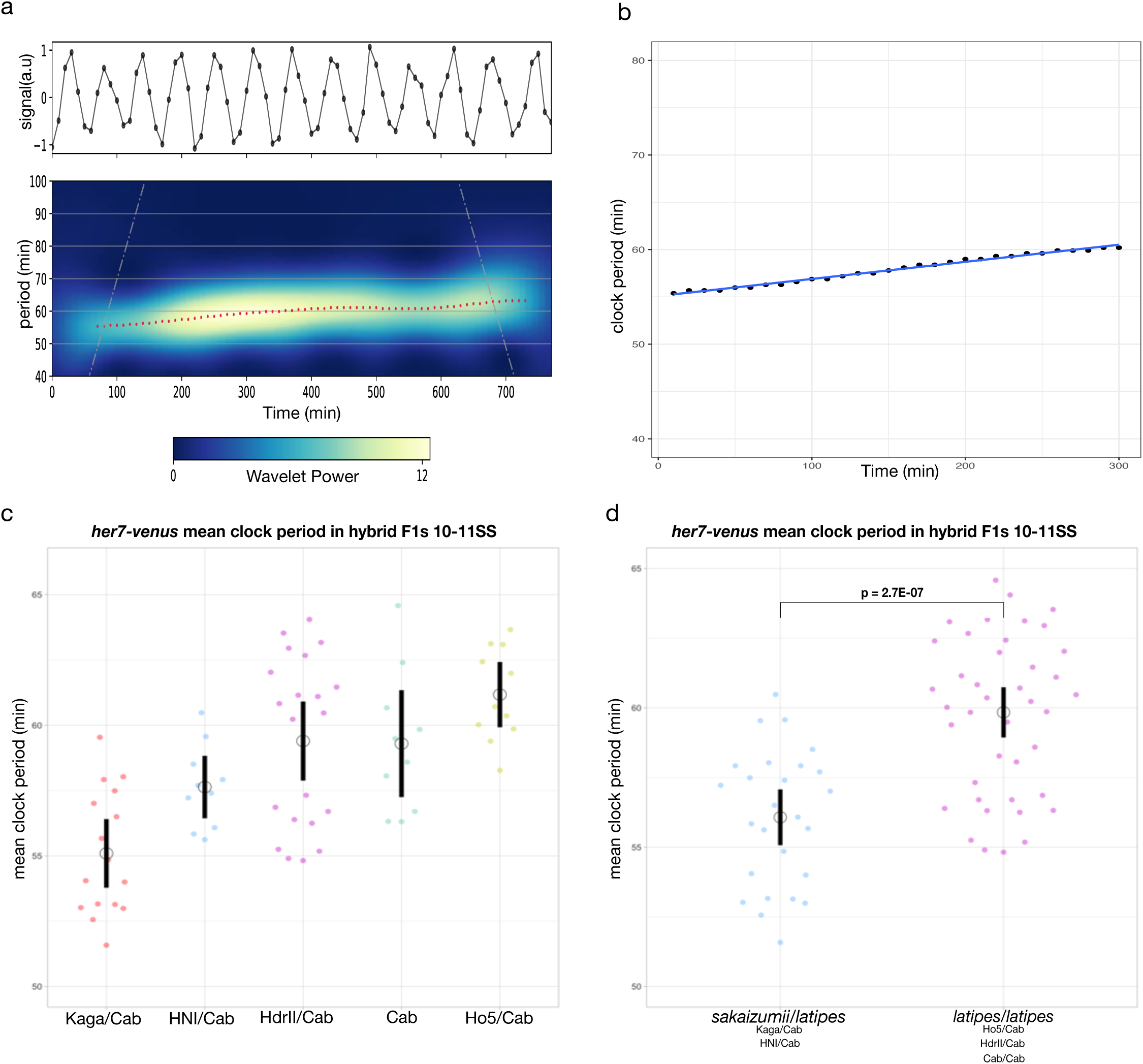
***her7-venus* period measurements in Hybrid F1 populations** (a) PyBoat wavelet transformation of detrended signal from Figure 2c shows continuous instantaneous period measurements over the course of live-imaging (b) example of period measurements every with 10 minute intervals for 300 minutes from the wavelet transformation in (a). Individual period measurements are black dots with linear-fit in blue. Mean period is the average of the black dot period measurements over 300 minutes, intercept period (clock period) is the y-intercept value of the blue fitted line. Period in minutes, Time in minutes. (c) endogenous *her7-venus* mean period measurements in hybrid F1 embryos at the 10-11SS. Kaga/Cab F1 hybrids have the fastest mean *her7-venus* period (55.09 minutes (sd +/- 2.39)), while HNI/Cab F1s show (57.63 minutes (sd +/- 1.59)), on the other hand hybrid F1 *O.latipes* strains Cab/Cab show (59.3 minutes (sd +/- 2.71)), Cab/HdrII show (59.4 minutes (sd +/- 3.24)), Cab/Ho5 show (61.17 minutes (sd +/-1,77)). Black circle = mean Black line = 95% confidence interval. One-way ANOVA p= 1.2E-06. Post-Hoc Tukey HSD testing shows significant differences between the following groups Kaga/Cab and Cab/Cab p adjusted = 1.2E- 03, Kaga/Cab and HdrII/Cab p adjusted = 4.0E-05, Kaga/Cab and Ho5/Cab p adjusted = 9.0E- 07, HNI/Cab and Ho5/Cab p adjusted = 2.1E-03. N= 16 Kaga/Cab F1, N= 10 HNI/Cab F1, N= 10 Cab/Cab F1, N= 21 HdrII/Cab F1, N= 11 Ho5/Cab F1. N= 16 Kaga/Cab F1, N= 10 HNI/Cab F1, N= 10 Cab/Cab F1, N= 21 HdrII/Cab F1, N= 11 Ho5/Cab F1. (d) endogenous *her7-venus* mean period measurements in hybrid F1 of *Oryzias sakaizumii/Oryzias latipes* or *Oryzias latipes/Oryzias latipes* embryos at the 10-11SS. *Oryzias sakaizumii/Oryzias latipes* F1 hybrids have a faster mean *her7-venus* period (56.07 minutes (sd +/- 2.43)) than *Oryzias latipes/Oryzias latipes* (59.84 minutes (sd +/- 2.86)). Black circle = mean Black line = 95% confidence interval. Welch two sample t-test p = 2.7E-07. N= 26 *Oryzias sakaizumii/Oryzias latipes* F1 hybrid embryos, N= 42 *Oryzias latipes/Oryzias latipes* F1 hybrid embryos.

**Extended Data Figure 3:**
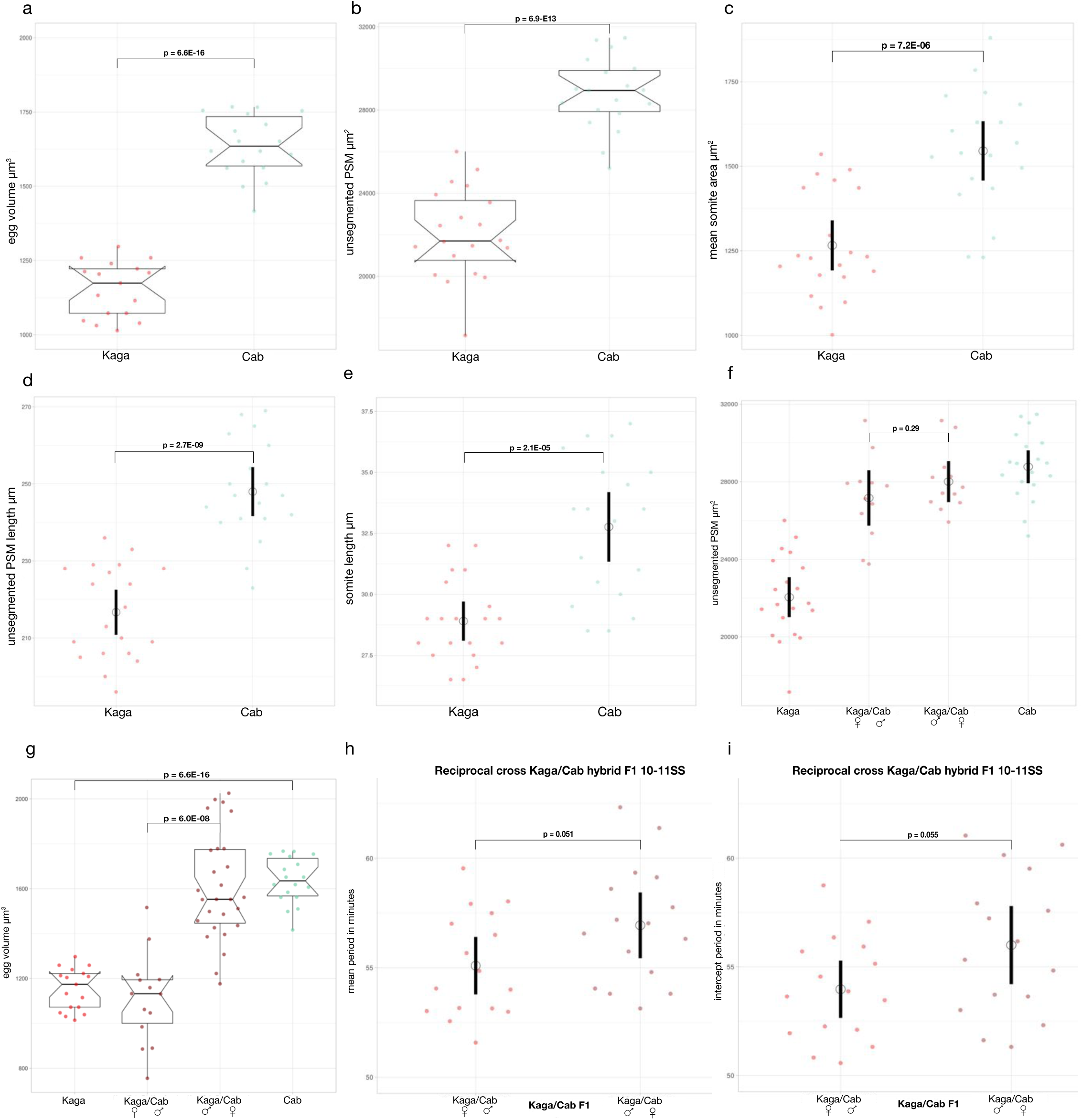
**Kaga and Cab F0 and reciprocal hybrid F1 segmentation timing and size measurements** (a) F0 Kaga have smaller fertilised egg volume (1152μm^3^ (sd +/- 93,0)) than Cab (1636μm^3^ (sd +/- 103)). Welch two sample t-test p = 6.6E-16. N= 17 Kaga eggs, N= 18 Cab eggs. (b) Area of unsegmented PSM at the 10-11 somite stage of Kaga and Cab F0 embryos. Kaga embryos have smaller unsegmented PSM (22048μm^2^ (sd +/- 2145)) compared to Cab (28770μm^2^ (sd +/- 1709)). Welch two sample t-test p = 6.9E-13. N= 20 Kaga embryos, N= 19 Cab embryos. (c) somite area of nascent pair of somites in Kaga and Cab F0 embryos at the 10-11 somite stage. Kaga embryos have smaller somites (1265μm^2^ (sd +/- 154)) than Cab (1545μm^2^ (sd +/- 177)) embryos. Black circle = mean Black line = 95% confidence interval. Welch two sample t-test p = 7.20E-06. N= 20 Kaga embryos, N= 19 Cab embryos. (d) length of unsegmented PSM at the 10SS of Kaga and Cab F0 embryos. Kaga embryos have shorter unsegmented PSM (216,7μm (sd +/- 12,19)) compared to Cab (248μm (sd +/- 12,83)). Black circle = mean Black line = 95% confidence interval. Welch two sample t-test p = 2.7E-09. N= 20 Kaga embryos, N= 19 Cab embryos. (e) length of nascent somite measured at the 10SS of Kaga and Cab F0 embryos. Kaga embryos make shorter somites (28.9μm (sd +/- 1.67)) compared to Cab (32.76μm (sd +/- 2.88)). Black circle = mean Black line = 95% confidence interval. Welch two sample t-test p = 2.1E-05. N= 20 Kaga embryos, N= 19 Cab embryos. (f) Area of unsegmented PSM μm^2^ at the 10-11SS of Kaga/Cab F1 reciprocal cross as compared to the paternal F0 Kaga and Cab populations. Kaga/Cab F1 embryos coming from Kaga females crossed to Cab male embryos have unsegmented PSM size of (2716μm^2^ (sd +/- 1592)) compared Kaga/Cab F1 embryos coming from Kaga males crossed to Cab females (28004μm^2^ (sd +/-2154)). While the Kaga F0 population embryos have an unsegmented PSM size of (22048μm^2^ (sd +/- 2145)) compared to Cab (28770μm^2^ (sd +/- 1709)). Black circle = mean Black line = 95% confidence interval. Welch two sample t-test p = 0.29. N= 20 Kaga embryos, N= 19 Cab embryos. N= 12 Kaga/Cab F1 embryos coming from a cross of Kaga females to Cab males, N= 12 Kaga/Cab F1 embryos coming from a cross of Kaga males to Cab females. (g) fertilised egg volume of Kaga/Cab F1 reciprocal cross shows a significant maternal effect. Kaga/Cab F1 embryos coming from Kaga females crossed to Cab male embryos have fertilised egg volume of (1110μm^3^ (sd +/- 198)) compared Kaga/Cab F1 embryos coming from Kaga males crossed to Cab females 1605μm^3^ (sd +/-236)). Welch two sample t-test p = 6.0E0-08. N= 14 Kaga/Cab F1 embryos coming from a cross of Kaga females to Cab males, N= 27 Kaga/Cab F1 embryos coming from a cross of Kaga males to Cab females. F0 Kagas have smaller fertilised egg volume (1152μm^3^ (sd +/- 93,0)) than Cab (1636μm^3^ (sd +/- 103)). Black circle = mean Black line = 95% confidence interval. Welch two sample t-test p = 6.6E-16. N= 17 Kaga eggs, N= 18 Cab eggs. (h) endogenous *her7-venus* mean period measurements in hybrid F1 Kaga/Cab reciprocal cross at the 10-11SS shows non-significant difference. Kaga/Cab F1 embryos coming from Kaga females crossed to Cab males have a mean *her7-venus* period of (55.09 minutes (sd +/- 2.39)) while Kaga/Cab F1 embryos coming from Kaga males crossed to Cab females have a mean period of (56.94 minutes (sd +/- 2.73)). Black circle = mean Black line = 95% confidence interval. Welch two sample t-test p = 0.051. N= 16 Kaga/Cab F1 embryos coming from a cross of Kaga females to Cab males, N= 16 Kaga/Cab F1 embryos coming from a cross of Kaga males to Cab females. (i) endogenous *her7-venus* intercept period measurements in hybrid F1 Kaga/Cab reciprocal cross at the 10-11SS shows non-significant difference. Kaga/Cab F1 embryos coming from Kaga females crossed to Cab males have an intercept *her7-venus* period of (53.97 minutes (sd +/- 2.40)) while Kaga/Cab F1 embryos coming from Kaga males crossed to Cab females have a mean period of (56.00 minutes (sd +/- 3.27)). Black circle = mean Black line = 95% confidence interval. Welch two sample t-test p = 0.055. N= 16 Kaga/Cab F1 embryos coming from a cross of Kaga females to Cab males, N= 16 Kaga/Cab F1 embryos coming from a cross of Kaga males to Cab females.

**Extended Data Figure 4:**
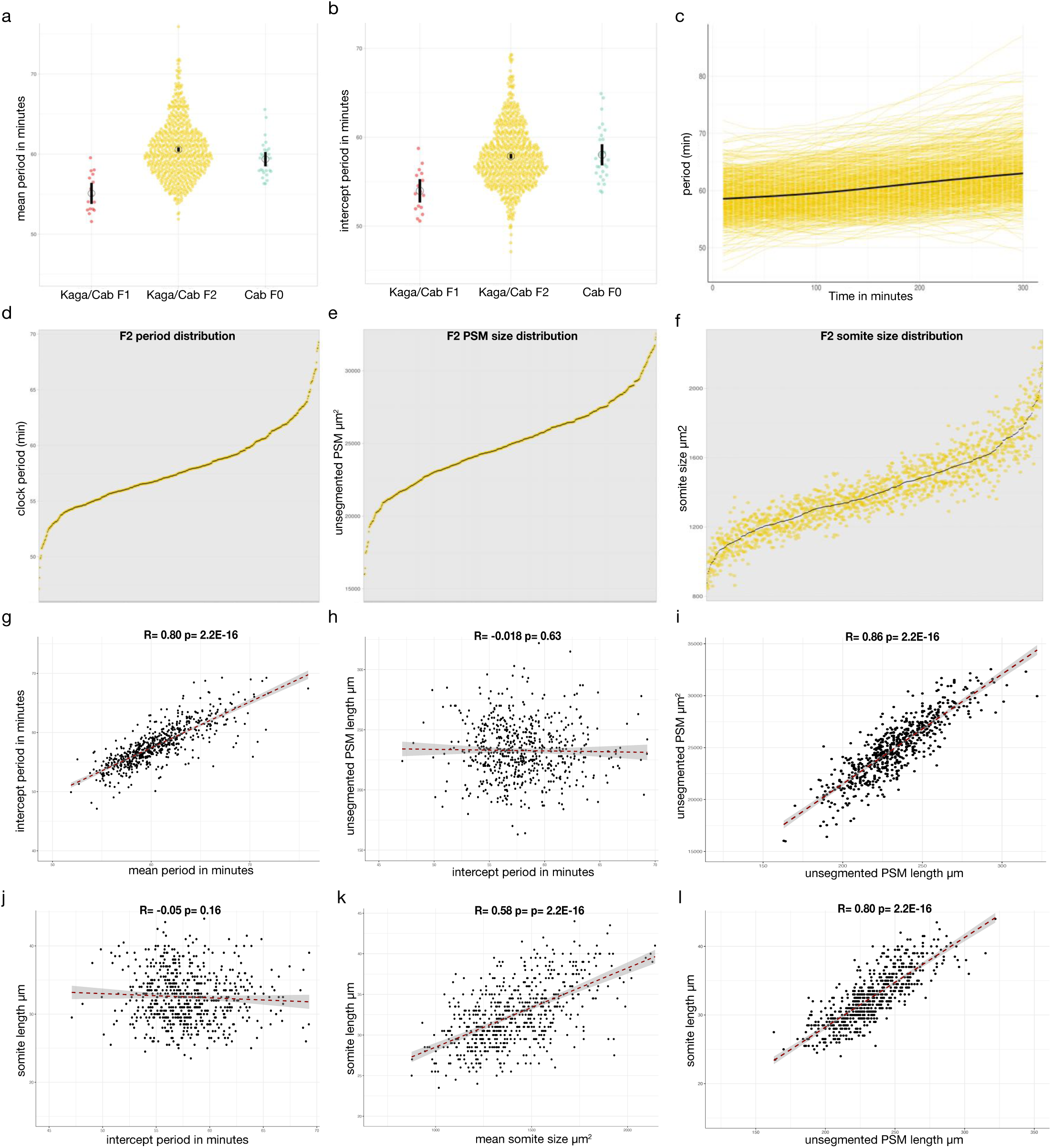
**F2 Kaga/Cab *her7-venus* reveals modular nature of segmentation timing and unsegmented PSM/somite size** (a) endogenous *her7-venus* mean period measurements of F2 Kaga/Cab embryos imaged at 10- 11SS (60.57 minutes (sd +/- 3.54)), compared to mean period values for F1 Kaga/Cab (55.09 minutes (sd +/- 2.39)) and F0 Cab/Cab (59.37 minutes (sd +/- 2.26)). Black circle = mean period. Black line = 95% confidence interval. Each dot is one embryo. N= 638 F2 Kaga/Cab embryos N= 16 F1 Kaga/Cab embryos N= 28 F0 Cab embryos. (b) endogenous *her7-venus* intercept period measurements of F2 Kaga/Cab embryos (57.86 minutes (sd +/- 3.43)) imaged at the 10-11SS compared to intercept period values for F1 Kaga/Cab (53.97 minutes (sd +/- 2.4)) and F0 Cab/Cab (58.03 minutes (sd +/- 3.02)). Black circle = mean period. Black line = 95% confidence interval. Each dot is one embryo. N= 638 F2 Kaga/Cab embryos N= 16 F1 Kaga/Cab embryos N= 28 F0 Cab embryos. Same panel as in Figure 3a (c) endogenous *her7-venus* period measurements over 300 minutes in F2 Kaga/Cab embryos imaged at the 10-11SS. A considerable endogenous *her7-venus* period range is obtained in the F2 cross. Black line = mean period N= 638 F2 Kaga/Cab embryos. (d) endogenous *her7-venus* clock period measurements of F2 Kaga/Cab arranged from fastest period (47.1 minutes) to slowest period (69.2 minutes). A significant period range (22.1 minutes) is obtained in the F2 cross. Each black line is an intercept period measurement from one F2 embryo. N= 638 F2 Kaga/Cab embryos. (e) Unsegmented PSM measurements of F2 Kaga/Cab arranged from smallest PSM 15986μm^2^ to the largest PSM 32546µm^2^. A significant PSM size range 16560µm^2^ is obtained in the F2 cross. Each black line is an unsegmented PSM measurement from one F2 embryo. N= 633 Kaga/Cab F2 (f) Somite size measurements are arranged from the smallest 875μm^2^ to the largest 2143μm^2^ . A sizable somite size range 1268μm^2^ is obtained in the F2 cross. Each black line is the mean somite size from one F2 embryo. Yellow dots represent individual (left and right) nascent somite size measurements. N= 631 Kaga/Cab F2. (g) Pearson’s correlation between mean period (a) and intercept period (b) measurements across all F2 Kaga/Cab embryos R= 0.80 p-value= 2.2E-16. Red dotted line= linear fit, grey shaded area= 95% confidence interval N= 638 F2 Kaga/Cab embryos (h) Pearson’s correlation between unsegmented PSM length and intercept period across all F2 Kaga/Cab embryos R= -=0.018 p-value= 63. Red dotted line= linear fit, grey shaded area= 95% confidence interval. N= 623 Kaga/Cab F2. (i) Pearson’s correlation between unsegmented PSM size and unsegmented PSM length across all F2 Kaga/Cab embryos R= 0.86 p-value= 2.2E-16. Red dotted line= linear fit, grey shaded area= 95% confidence interval. N= 632 Kaga/Cab F2. (j) Pearson’s correlation between somite length and intercept period in minutes across all F2 Kaga/Cab embryos R= - 0.05 p-value= 0.16. Red dotted line= linear fit, grey shaded area= 95% confidence interval. N= 623 Kaga/Cab F2. (k) Pearson’s correlation between somite length and mean somite size across all F2 Kaga/Cab embryos R= 0.58 p-value= 2.2E-16. Red dotted line= linear fit, grey shaded area= 95% confidence interval. N= 632 Kaga/Cab F2. (l) Pearson’s correlation between somite length and unsegmented PSM length across all F2 Kaga/Cab embryos R= 0.80 p-value= 2.2E- 16. Red dotted line= linear fit, grey shaded area= 95% confidence interval N= 632 Kaga/Cab F2.

**Extended Data Figure 5:**
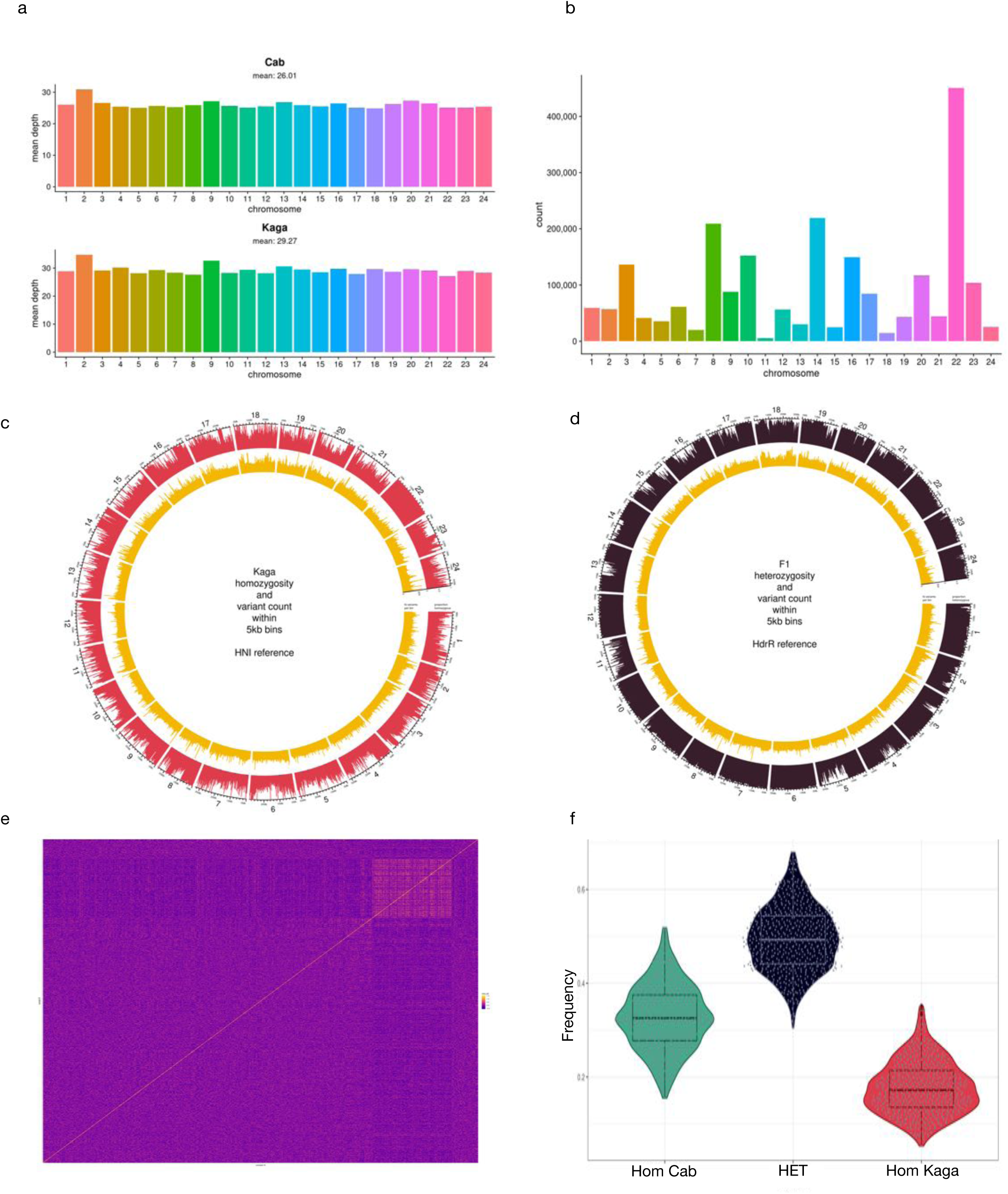
**Whole genome sequencing of F0, F1 and F2 Kaga/Cab** (a) Mean sequencing depth per chromosome for *Cab* and *Kaga* F0 strains, with genome-wide mean depth across all chromosomes shown (b) Number of SNPs per chromosome that are homozygous-divergent between F0 *Cab* and *Kaga*, and heterozygous in the F1 generation.(c) Circos plot showing the 24 chromosomes of a whole genome sequenced F0 Kaga embryo aligned against the HNI northern reference genome. Proportion of homozygous SNPs within 5kb bins in the Kaga F0 genome is shown in red and number of SNPs in each bin in yellow. The mean homozygoisty across all bins is 31% indicating a moderate level of isogenecity genome-wide (d) Circos plot showing the 24 chromosomes of a whole genome sequenced F1 hybrid Kaga/Cab embryo aligned against the HdrII reference genome. Proportion of heterozygous SNPs within 5kb bins in the F1 hybrid genome is shown in dark brown and number of SNPs in each bin in yellow. The mean heterozygosity across all bins is 67% (e) A genetic relationship matrix (GRM) constructed from the entire genome of 600 F2 samples is presented as a heatmap, with each sample represented on each axis, and lighter colours representing a higher degree of relatedness between a pair of samples. The square in the top right-hand corner is created by samples 550-648, which have distinct genotypes to the rest of the samples due to their having been bred from different F1 parents. (f)Proportions of 5-kb blocks called as either homozygous-*Cab*, heterozygous, or homozygous-*Kaga* in the F2 Kaga/Cab embryos.

**Extended Data Figure 6:**
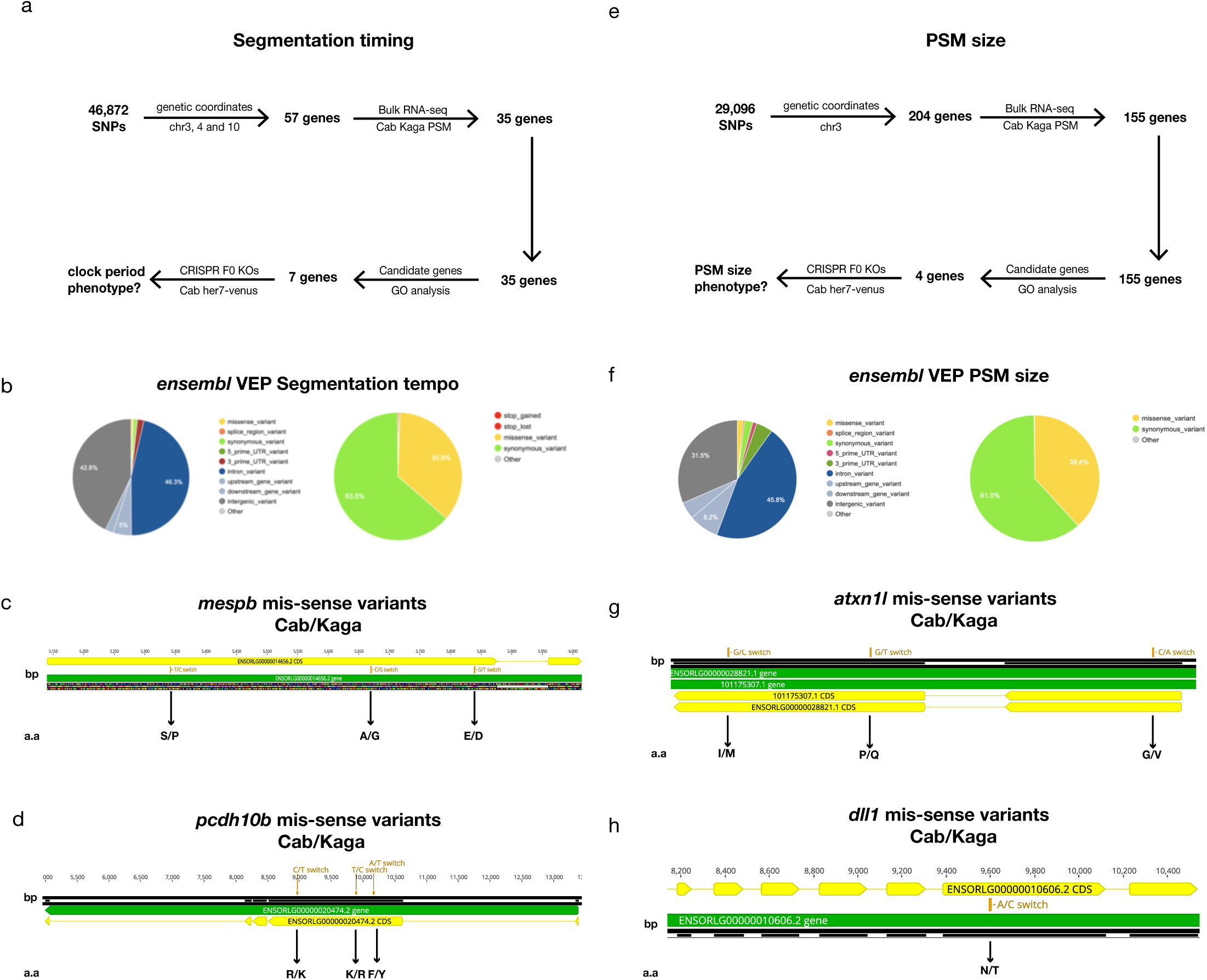
***dev*QTL mapping and annotation for segmentation timing and size** (a) Workflow from *dev*QTL mapping to candidate gene selection. For segmentation timing 46,872 homozygous divergent SNPs between Kaga and Cab were located on chromosomes 3, 4 and 10. Genomic coordinates revealed a total of 57 genes on all 3 chromosomes located in regions that passed the significance threshold. Bulk RNA-seq on Kaga and Cab unsegmented PSM narrowed down the number of candidate genes expressed in the PSM to 35 genes. GO enrichment and candidate gene picking led to a top 7 gene list which were selected to perform F0 CRISPR KOs in *her7-venus* Cab background to check for a clock period phenotype (b) *ensembl* variant effect predictor (VEP) output for segmentation timing showing the distribution of homozygous divergent SNPs between Kaga and Cab in the F2 Kaga/Cab QTL mapping on segmentation timing, most of the divergent SNPs fall in either intronic or intergenic regions, of the ones that fall within the coding sequence of genes the majority lead to synonymous mutations (63.5%) while only a minority (35.9%) lead to miss-sense mutations (c) position of mis-sense variants between Cab/Kaga in the coding sequence of *mespb.* Three base-pair (bp) changes cause 3 amino acid (a.a) changes: Serine/Proline (S/P), Alanine/Glycine (A/G) and Glutamic Acid/Aspartic Acid (E/D) are highlighted. Visualisation done using *Geneious*. (d) position of mis-sense variants between Cab/Kaga in the coding sequence of *pcdh10b.* Three base-pair (bp) changes cause 3 amino acid (a.a) changes: Arginine/Lysine (R/K), Lysine/Arginine (K/R) and Phenylalanine/Tyrosine (F/Y) are highlighted. Visualisation using *Geneious*. (e) Workflow from *dev*QTL mapping to candidate gene selection. For PSM size 29,096 homozygous divergent SNPs between Kaga and Cab are located on chromosomes 3. Genomic coordinates revealed a total of 204 genes located in regions that passed the significance threshold. Bulk RNA-seq on Kaga and Cab unsegmented PSM narrowed down the number of candidate genes expressed in the PSM to 155 genes. GO enrichment and candidate gene picking led to a top 4 genes which were selected to perform F0 CRISPR KOs in Cab background to check for PSM size phenotype. (f) VEP output for PSM size showing the distribution of homozygous divergent SNPs between Kaga and Cab in the F2 Kaga/Cab QTL on PSM size, majority of the divergent SNPs fall in either intronic or intergenic regions, of the ones that fall within the coding sequence of genes the majority lead to synonymous mutations (61.5%) while only a minority (38.4%) lead to miss-sense mutations. (g) position of mis-sense variants between Cab/Kaga in the coding sequence of *atxn1l.* Three base-pair (bp) changes cause 3 amino acid (a.a) changes: Isoleucine/Methionine (I/M), Proline/Glutamine (P/Q) and Glycine/Valine (G/V) are highlighted. Visualization using *Geneious.* (h) position of mis-sense variants between Cab/Kaga in the coding sequence of *dll1.* One base-pair (bp) change causes 1 amino acid (a.a) change: Asparagine/Threonine (N/T) is highlighted. Visualisation using *Geneious*.

**Extended Data Figure 7:**
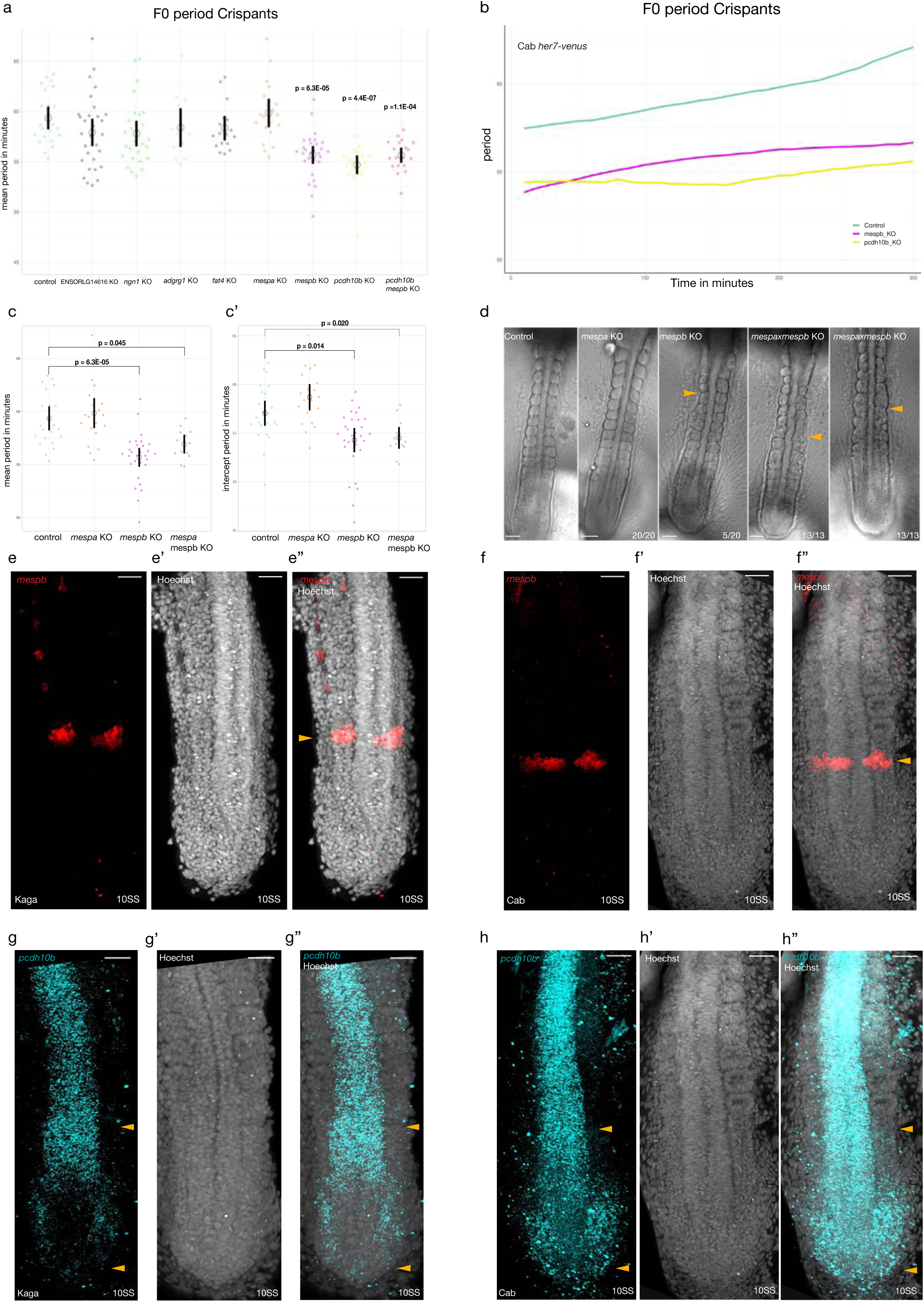
**Functional validation of *dev*QTLs on segmentation timing using F0 Crispants and HCR** (a) endogenous *her7-venus* mean period measurements in Control, *ENSORLG14616, ngn1, adgrg1, fat4, mespa, mespb*, *pcdh10b, mespb*+*pcdh10b* F0 Cab Crispants imaged at the 10- 11SS. Kruskal-Wallis’ test p= 7.8E-11. Post-Hoc Dunn’s test: only *mespb* (p-adjusted=6.3E- 05), *pcdh10b* (p-adjusted =4.4E-07)*, mespb*+*pcdh10b* (p-adjusted=1.1E-04) show a significant difference in mean period as compared to control injected embryos. N= 23 control injected Cab *her7-venus* embryos, N= 30 *ENSORLG14616,* N= 27 *ngn1,* N= 13 *adgrg1,* N= 17 *fat4,* N= 20 *mespa,* N= 29 *mespb,* N= 20 *pcdh10b,* N= 19 *pcdh10b+mespb* CRISPR/Cas9 injected into Cab *her7-venus.* (b) endogenous *her7-venus* posterior period measurements over 300 minutes in 10-11SS Cab F0 CRISPR/Cas9 knock-outs on candidate genes from the *dev*QTL. *mespb* (magenta) and *pcdh10b* (yellow) F0 Crispants have a faster segmentation period than Control Cas9 mRNA injected *her7-venus* Cab embryos (green). Period lines= mean values across samples. N= 23 control injected Cab *her7-venus* embryos. N= 29 *mespb* CRISPR/Cas9 injected into Cab *her7-venus.* N= 20 *pcdh10b* CRISPR/Cas9 injected into Cab *her7-venus.* (c) endogenous *her7-venus* mean period measurements in Control, *mespa*, *mespb* and *mespa+mespb* F0 Cab Crispants imaged at the 10-11SS. *mespb* and *mespa+mespb* Crispants show a faster mean *her7-venus* period (55.67 minutes (sd +/- 2.3)) and (56.92 minutes (sd +/- 1.41)) respectively than Cab control Cas9 mRNA injected embryos (59.35 minutes (sd +/- 2.56)) and *mespa* Crispants (59.85 minutes (sd +/- 2.96)). Kruskal-Wallis’ test p= 7.8E-11. Post-Hoc Dunn’s test: only *mespb* (p-adjusted=6.3E-05)*, mespa*+*mespb* (p-adjusted=0.045) show a significant difference in mean period as compared to control injected embryos. N= 23 control injected Cab *her7-venus* embryos, N= 29 *mespb,* N= 20 *mespa,* N= 13 *mespa+mespb* CRISPR/Cas9 injected into Cab *her7-venus.* (c’) endogenous *her7-venus* intercept period measurements in Control, *mespa*, *mespb* and *mespa+mespb* F0 Cab Crispants imaged at the 10-11SS. *mespb* and *mespa+mespb* Crispants show a faster intercept *her7-venus* period (54.28 minutes (sd +/- 3.2)) and (54.51 minutes (sd +/- 1.8)) respectively than Cab control Cas9 mRNA injected embryos (57,04 minutes (sd +/- 2.9)) and *mespa* Crispants (58.69 minutes (sd +/- 2.81)) Kruskal-Wallis’ test p= 8.9E-08. Post-Hoc Dunn’s test: only *mespb* (p-adjusted=0.014)*, mespa*+*mespb* (p-adjusted=0.020) show a significant difference in intercept period as compared to control injected embryos. N= 23 control injected Cab *her7-venus* embryos, N= 20 *mespa,* N= 29 *mespb,* N= 13 *mespa+mespb* CRISPR/Cas9 injected into Cab *her7-venus.*(d) Brightfield imaging of tails of Control, *mespa*, *mespb* and *mespa+mespb* F0 Cab Crispants imaged at the 10-11SS. *mespa* Crispants (20/20) show no morphological somite phenotype, while a minority *mespb* Crispants (5/20) show mild defects in one or more somite boundaries and size (yellow arrowheads). All *mespa+mespb* (13/13) double Crispants show severe defects in somite boundary formation and somite size (yellow arrowheads). Scale bar= 50µm. (e-e’’) HCR on *mespb* performed on Kaga embryos at the 10SS. (e) *mespb* is expressed as one stripe at the S-1 somite position (e’) Hoechst labels all nuclei in the tail (e’’) overlay of *mespb* and Hoechst from (e-e’), yellow arrowhead highlights position of the *mespb* stripe in relation to the forming somites in the tail. N= 8 embryos, Scale bar= 30µm. (f-f’’) HCR on *mespb* performed on Cab embryos at the 10SS. (f) *mespb* is expressed as one stripe at the S-1 somite position (f’) Hoechst labels all nuclei in the tail (f’’) overlay of *mespb* and Hoechst from (f-f’), yellow arrowhead highlights position of the *mespb* stripe in relation to the forming somites in the tail. N= 12 embryos, Scale bar= 40µm. (g-g’’) HCR on *pcdh10b* performed on Kaga embryos at the 10SS. (g) *pcdh10b* is expressed in the neural tube in addition to the unsegmented PSM. Anterior and posterior PSM positions are highlighted by yellow arrowheads (g’) Hoechst labels all nuclei in the tail (g’’) overlay of *pcdh10b* and Hoechst from (g-g’), yellow arrowhead highlights position of anterior and posterior PSM. N= 9 embryos, Scale bar= 30µm. (h-h’’) HCR on *pcdh10b* performed on Cab embryos at the 10SS. (h) *pcdh10b* is expressed in the neural tube in addition to the unsegmented PSM. Anterior and posterior PSM positions are highlighted by yellow arrowheads, higher expression levels in the PSM domain and an extended expression in the neural tube are observed as compared to Kaga embryos (h’) Hoechst labels all nuclei in the tail (h’’) overlay of *pcdh10b* and Hoechst from (g-g’), yellow arrowhead highlights position of anterior and posterior PSM. N= 8 embryos, Scale bar= 40µm.

**Extended Data Figure 8:**
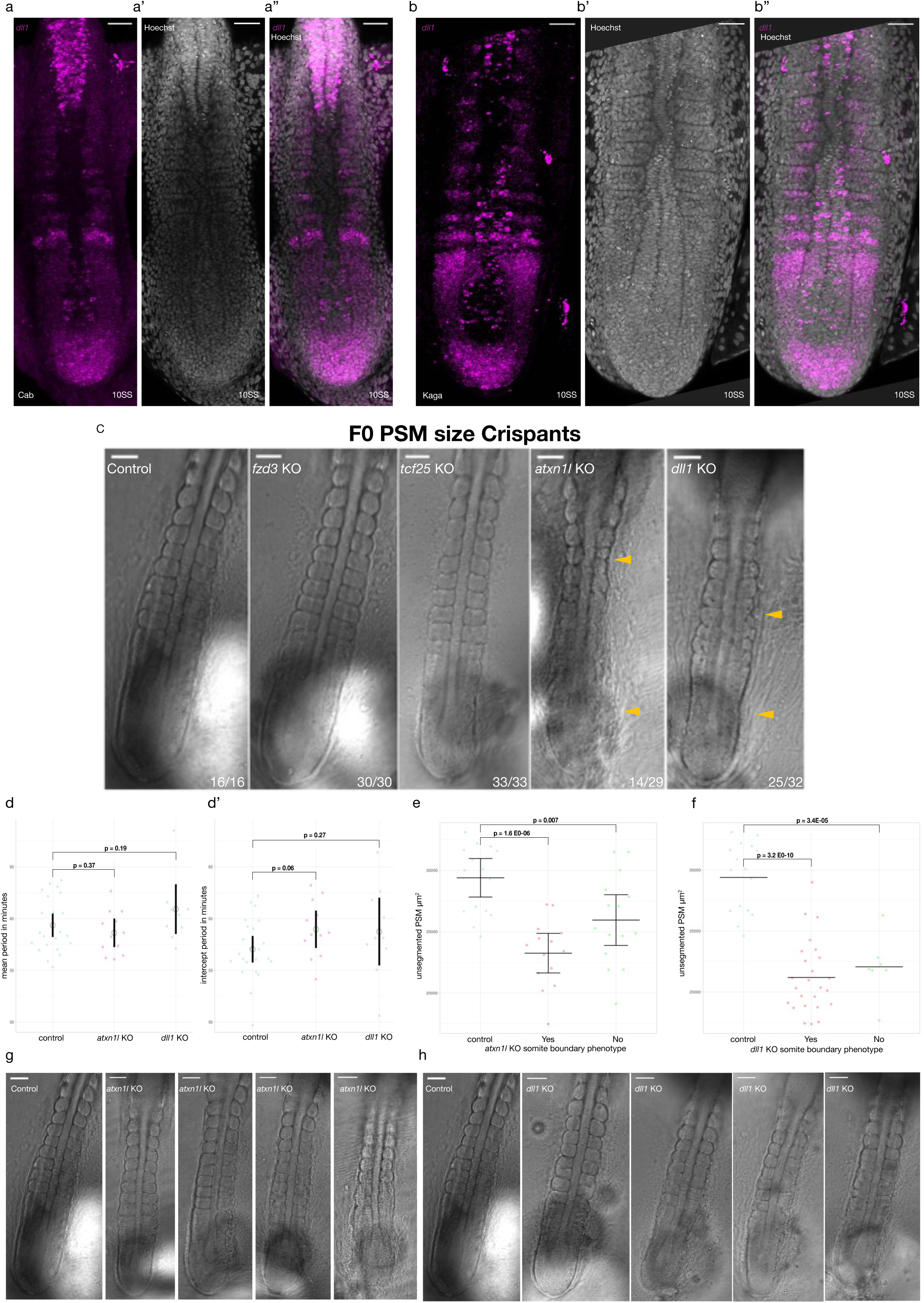
**Functional validation of *dev*QTLs on unsegmented PSM size using F0 Crispants and HCR** (a-a’’) HCR on *dll1* performed on Cab embryos at the 10SS. (a) *dll1* is expressed throughout the unsegmented PSM (posterior and anterior), in addition to striped expression in formed somites (a’) Hoechst labels all nuclei in the tail (a’’) overlay of *dll1* and Hoechst from (a-a’). N= 6 embryos, Scale bar= 40µm. (b-b’’) HCR on *dll1* performed on Kaga embryos at the 10SS. (b) *dll1* is expressed throughout the unsegmented PSM (posterior and anterior), in addition to striped expression in formed somites (b’) Hoechst labels all nuclei in the tail (b’’) overlay of *dll1* and Hoechst from (b-b’). N= 7 embryos, Scale bar= 40µm. (c) unsegmented PSM size Cab F0 CRISPR/Cas9 knock-outs performed on candidate genes from the *dev*QTL mapping results, brightfield imaging was carried out on Control, *fzd3*, *tcf25, atxn1l* and *dll1* F0 Cab Crispants imaged at the 10-11 somite stage. Both *atxn1l* and *dll1* Crispants showed smaller unsegmented PSM sizes than control injected embryos (yellow arrow-heads), in addition both *atxn1l* (14/29) and *dll1* (25/32) Crispants showed morphological somite size and boundary abnormalities (yellow arrow-heads). N= 16 control injected embryos,N = 30 *fzd3* Crispants N= 33 *tcf25* Crispants N= 29 *atxn1l* Crispants N= 32 *dll1* Crispants. Scale bar= 50µm.(d) endogenous *her7-venus* mean period measurements in Control, *atxn1l* and *dll1* F0 Cab Crispants imaged at the 10-11 somite stage. Both Crispants show a similar mean *her7-venus* period (58.63 minutes (sd +/- 2.08)) and (60.92 minutes (sd +/- 3.18)) respectively to Cab control Cas9 mRNA injected embryos (59.35 minutes (sd +/- 2.56)). Welch two sample t-test p = 0,37 for *atxn1l* Crispants and p = 0,19 for *dll1* Crispants. N= 23 control injected Cab *her7-venus* embryos. N= 12 *atxn1l,* N= 10 *dll1* CRISPR/Cas9 F0 injected into Cab *her7-venus.* (d’) endogenous *her7-venus* intercept period measurements in Control, *atxn1l* and *dll1* F0 Cab Crispants imaged at the 10- 11 somite stage. Both Crispants show a similar intercept *her7-venus* period (58.97 minutes (sd +/- 2.73)) and (58.75 minutes (sd +/- 4.34)) respectively to Cab control Cas9 mRNA injected embryos (57.04 minutes (sd +/- 2.90)). Welch two sample t-test p = 0.064 for *atxn1l* Crispants and p = 0.27 for *dll1* Crispants. N= 23 control injected Cab *her7-venus* embryos. N= 12 *atxn1l,* N= 10 *dll1* CRISPR/Cas9 F0 injected into Cab *her7-venus.* (e) unsegmented PSM size measurements on 10-11 somite stage Cab F0 CRISPR/Cas9 knock-outs separated by the presence or absence of a somite morphology phenotype in *atxn1l* Crispants. *atxn1l* Cripsants with no somite phenotype still showed significantly smaller unsegmented PSM size (25947µm^2^ (sd +/- 3605)) when compared to Cab control Cas9 mRNA injected embryos (29392µm^2^ (sd +/- 2851)). *atxn1l* Cripsants with a somite phenotype also showed significantly smaller unsegmented PSM size (23251µm^2^ (sd +/- 2689)) compared to Control injected samples. Welch two sample t-test p = 7.1E-03 for *atxn1l* Crispants with no somite phenotype and p = 1,5E-06 for *atxn1l* Crispants with a somite phenotype. N = 16 control injected embryos N= 15 *atxn1l* Crispants with no somite phenotype N= 14 *atxn1l* Crispants with visible somite phenotype (f) unsegmented PSM size measurements on 10-11SS Cab F0 CRISPR/Cas9 knock-outs separated by the presence or absence of a somite phenotype in *dll1* Crispants. *dll1* Cripsants with no somite phenotype (minority of samples) showed significantly smaller unsegmented PSM size (22064µm^2^ (sd +/- 2513)) when compared to Cab control Cas9 mRNA injected embryos (29392µm^2^ (sd +/- 2851)). *dll1* Cripsants with a somite phenotype showed significantly smaller unsegmented PSM size (21191µm^2^ (sd +/- 2977)) compared to Control injected samples. Welch two sample t-test p = 3.4E-05 for *dll1* Crispants with no somite phenotype and p = 3.19E-10 for *dll1* Crispants with a somite phenotype. N = 16 control injected embryos N= 7 *dll1* Crispants with no somite phenotype N= 25 *dll1* Crispants with visible somite phenotype (g) Brightfield imaging on control injected embryos and *atxn1l* F0 Cab Crispants imaged at the 10-11SS showing no somite phenotype. Example of 3 *atxn1l* Crispants with smaller unsegmented PSM sizes and no visible somite phenotypes compared to control injected embryos. N= 16 control injected embryos, N = 15 *atxn1l* Crispants with no somite morphology phenotype. Scale bar= 50µm. (h) Brightfield imaging on control injected embryos and *dll1* F0 Cab Crispants imaged at the 10-11SS showing no visible somite phenotype. Example of 3 *dll1* Crispants with smaller unsegmented PSM sizes and no somite morphology phenotype compared to control injected embryos (same embryo as in H). N= 16 control injected embryos, N= 7 *dll1* Crispants with no somite morphology phenotype. Scale bar= 50µm.

## Supplementary Information

**Supplementary Information 1:** custom-made Fiji macro-script for concatenating and extracting fluorescent intensity measurements from a defined ROI

**Supplementary Information 2:** gene-counts table for bulk RNA-seq on unsegmented PSM of Cab and Kaga tails

**Supplementary Information 3:** Ensembl ID and sequences of gRNAs used

## Notes

### Competing Interest Statement

The authors have declared no competing interest.

