## Supplementary material for "Modular control of time and space during vertebrate axis segmentation": SI_1

path0 = File.openDialog("Select the BF image path");

dir0 = File.getParent(path0);

name0 = File.getName(path0);

basename = substring(name0,0,lengthOf(name0)-5);

for (l=1; l<=11; l++) {

      j = l;

      print(j);

if (l==10){

basename = substring(basename,0,lengthOf(basename)-1);

}

open(dir0 +"/"+basename+l+".tif");

roiManager("Open", dir0 +"/"+basename+l+".zip");

roiManager("Sort");

roiManager("List");

selectWindow("Overlay Elements of "+basename+l+".tif");

table = getInfo("window.contents");

ROIs = split(table, "\n");

ROIline=split(ROIs[ROIs.length-1], "\t");

time=(ROIline[14]);

diameter=(ROIline[6]);

XArray=newArray(time);

YArray=newArray(time);

for (i=1; i<(ROIs.length-1); i++) {

ROIline1=split(ROIs[i], "\t");

x1U=(ROIline1[4]);

y1U=(ROIline1[5]);

t1U=(ROIline1[14]);

XArray[(t1U-1)]=x1U;

YArray[(t1U-1)]=y1U;

ROIline2=split(ROIs[i+1], "\t");

x2U=(ROIline2[4]);

y2U=(ROIline2[5]);

t2U=(ROIline2[14]);

dx=(x2U-x1U)/(t2U-t1U);

dy=(y2U-y1U)/(t2U-t1U);

for (k=1; k<(t2U-t1U); k++) {

XArray[(t1U-1+k)]=x1U+(k\*dx);

YArray[(t1U-1+k)]=y1U+(k\*dy);

}

}

ROIline1=split(ROIs[ROIs.length-1], "\t");

xLU=(ROIline1[4]);

yLU=(ROIline1[5]);

tLU=(ROIline1[14]);

XArray[(tLU-1)]=xLU;

YArray[(tLU-1)]=yLU;

//roiManager("Save", dir0+"/"+substring(name0,0,lengthOf(name0)-4)+".zip");

selectWindow(basename+l+".tif");

Stack.setPosition(1, 1, 1); //(channel, slice, frame)

k = newArray();

//loop over frames

for (m=1; m<=tLU; m++) {

Stack.setFrame(m);

//make selection circle

run("Overlay Options...", "stroke=cyan width=1 fill=none set");

makeOval(XArray[m-1],YArray[m-1],diameter,diameter);

run("Measure");

k = Array.concat(k,getResult("Mean"));

run("Add Selection...");

}

run("Clear Results");

for (j=0; j<k.length; j++) {

  setResult("Value", j, k[j]); }

updateResults();

saveAs("Measurements",dir0 +"/"+basename+l+".csv");

roiManager("Deselect");

roiManager("Delete");

selectWindow("Overlay Elements of "+basename+l+".tif");

run("Close");

selectWindow("Results");

run("Close");

saveAs("Tiff", dir0+"/overlayed/"+name0);

close();

}
